# Quantitative Comparison of Presenilin Protein Expression Reveals Greater Activity of PS2-γ-Secretase

**DOI:** 10.1101/2023.05.09.540102

**Authors:** Melissa Eccles, Nathan Main, Miheer Sabale, Brigid Roberts-Mok, Mark Agostino, David Groth, Paul E. Fraser, Giuseppe Verdile

**Affiliations:** Curtin Medical School, Curtin Health Innovation Research Institute (CHIRI), Curtin University, Bentley, Western Australia, Australia; Dementia Research Centre, Macquarie Medical School, Faculty of Medicine, Health and Human Sciences, Macquarie University, Sydney, New South Wales, Australia; Tanz Centre for Research in Neurodegenerative Diseases, University of Toronto, Toronto, Ontario M5T 2S8; Department of Medical Biophysics, University of Toronto; School of Medical and Health Sciences, Edith Cowan University, Joondalup, Western Australia, Australia

**Keywords:** Presenilin-1, Presenilin-2, γ-secretase, APP, Notch1, amyloid-β, Quantitative Presenilin Expression

## Abstract

γ-Secretase processing of APP has long been of interest in the pathological progression of Alzheimer’s disease (AD) due to its role in the generation of amyloid-β. The catalytic component of the enzyme are the presenilins of which there are two homologues, Presenilin-1 (PS1) and Presenilin-2 (PS2). The field has focussed on the PS1 form of this enzyme, as it is typically considered the more active at APP processing. However, much of this work has been completed without appropriate consideration of the specific levels of protein expression of PS1 and PS2. We propose that expression is an important factor in PS1- and PS2-γ-secretase activity, and that when this is considered, PS1 does not have greater activity than PS2. We developed and validated tools for quantitative assessment of PS1 and PS2 protein expression levels to enable direct comparison of PS protein in exogenous and endogenous expression systems, in HEK-293 PS1 and/or PS2 knockout cells. We show that exogenous expression of Myc-PS1-NTF is 5.5-times higher than and Myc-PS2-NTF. Quantitating endogenous PS protein levels using a novel PS1/2 fusion standard we developed showed similar results. When the marked difference in PS1 and PS2 protein levels is considered, we show that compared to PS1-γ-secretase, PS2-γ-secretase has equal or more activity on APP and Notch1. This study has implications for understanding the PS1 and PS2 specific contributions to substrate processing, and their potential influence in AD pathogenesis.

## INTRODUCTION

Presenilin (PS) is the catalytic component of γ-secretase, a tetrameric enzyme that cleaves type I transmembrane proteins. The two PS homologues, PS1 and PS2, share approximately 67% amino acid sequence similarity, and form active γ-secretase complexes when incorporated with Nicastrin, Anterior Pharynx Defective-1, and Presenilin Enhancer-2.^1^ The γ-secretase enzyme has been shown to cleave a large repertoire of substrates,^2^ the most well investigated of which are Amyloid Precursor Protein (APP)^3^ and Notch1.^4^ The cleavage of APP has received the most attention as it ultimately results in the generation of amyloid-β (Aβ) peptides, accumulation of which contributes to Alzheimer’s disease (AD) pathogenesis.^5^ Consequently, γ-secretase has been proposed as a therapeutic target, with the development of inhibitors of PS related γ-secretase activity.^6–10^ However, these molecules have failed in clinical trials due to off-target effects, which are thought to be caused by the inhibition of substrates other than APP, particularly Notch1,^7, 11, 12^ and may be influenced by differences in affinity for PS1- and PS2-γ-secretase.^13–15^ More recently the focus has shifted to the development of γ-secretase modulators (GSMs), small molecules that modulate the type of Aβ peptides released, while still maintaining cleavage of the intracellular domain of substrates.^16–18^ However, there is still a need for improved understanding of γ-secretase activity and insight into the differing roles of PS1- and PS2-γ-secretase enzymes.

γ-Secretase cleaves its substrates via a process termed **R**egulated **I**ntramembrane **P**roteolysis (RIP), where type I transmembrane proteins undergo multiple cleavages as part of the signalling or degradation processes. The first step in RIP is the shedding of the substrate ectodomain by proteases (in particular, ADAM (a disintegrin and metalloproteinase) family enzymes and aspartyl proteases BACE1 (β-APP cleaving enzyme) and BACE2.^19, 20^ The second step is performed by γ-secretase, where multiple intramembrane cleavages of the substrate transmembrane domain lead to release of the **I**ntra**C**ellular **D**omain (ICD) and secreted peptides.^3, 21^ In the case of APP, RIP can be initiated by either ADAM or BACE cleavage,^22, 23^ it is BACE cleavage that initiates the amyloidogenic pathway leading to the generation of Aβ peptides. γ-Secretase is known to successively ‘trim’ APP after the initial cleavage by tri- and tetrapeptide cleavages until the Aβ peptide is released from the luminal membrane.^24, 25^ It must be acknowledged that much of how APP is cleaved has been determined via investigations of PS1-γ-secretase, with little understanding of whether this process differs for PS2-γ-secretase.

The focus on PS1 appears to be largely a result of the significantly greater number of familial AD causing missense mutations in *PSEN1* (200+) compared to *PSEN2* (20+) (retrieved from www.alzforum.org/mutations September 2022).^26^ While both *PSEN1* and *PSEN2* mutations generally cause increased Aβ42:40 ratios, *PSEN1* mutations have an earlier average age of onset and are typically more aggressive.^27^ However, the recent identification of a *PSEN2* variant in the 3’ UTR that mutates a miRNA-binding region suggests PS2 protein expression may influence AD pathology.^28, 29^ This mutation has been shown to cause upregulated PS2 protein expression and subsequently increased Aβ42/Aβ40 ratio.^28^

While there is considerable functional overlap between PS1- and PS2-γ-secretase, there are several differences. Subcellular localisation has been shown to differ, with PS2-γ-secretase localised to late endosomal and lysosomal compartments,^30–32^ while the localisation of PS1-γ-secretase predominately resides within the plasma membrane.^31, 32^ As ectodomain shedding by BACE is a prerequisite for Aβ formation, its localisation in intracellular organelles, including endosomal compartments,^33^ links the PS2-γ-secretase complex to Aβ generation. PS2 has been shown to generate significantly more intracellular Aβ^32, 34^ and produce a higher Aβ42:Aβ40 ratio,^30, 35–37^ supporting its greater activity within the endosomal-lysosomal cellular compartment.

One aspect of PS1- and PS2-γ-secretase activity that is not often considered is expression within cells and tissues. Evidence from post-mortem tissues and in-vivo studies suggest that PS expression levels vary with age and other AD-associated changes. Lee et al,^38^ show that transcript expression of PS1 is significantly higher than PS2 in human foetal cortex, and that following birth and with age, a concomitant decrease in PS1 and increase in PS2 leads to approximately equal PS1 and PS2 expression. A similar PS expression profile has been observed during terminal differentiation of iPSC-derived neurons, where PS1 expression decreases and PS2 expression increases.^30^ Interestingly, PS1 protein expression is decreased in human AD cortex and hippocampus,^39^ and in aged murine cortex, there is a concomitant decrease in PS1 and increase in PS2 protein expression.^40^ These observations suggest a role for PS2 in neuronal maturation and, together with the role of PS2-γ-secretase in generating intracellular Aβ and increased Aβ42 product, may indicate that PS2 contributes more to AD pathology than previously credited.

In-vitro studies comparing PS1- and PS2-γ-secretase activity have typically shown PS1-complexes to be more active at processing APP and Notch. However, a major limitation of this work, is the assumption that PS1 and PS2 protein expression levels are equal. The comparable assessment of PS1 and PS2 expression is difficult, as there is no common PS antibody that detects both PS1 and PS2. To address this inability to directly assess PS1 and PS2 endogenous expression, activity is often assessed in cells where both PS1 and PS2 have been ablated from the cells and the PS is exogenously re-introduced to the cell. This presents an opportunity to tag the exogenous PS enabling equi-detection. However, to our knowledge, this approach has only been presented twice; firstly for determining the cellular localisation of PS1 vs. PS2,^32^ and secondly for use in PS quantitation, after which it was determined that, when PS expression was considered, there was no significant difference in γ-secretase activity.^41^ Other studies using the relative levels of mature Nct to normalise for exogenous PS expression have shown discordant results; no difference in APP and Notch ICD generation,^42^ or reduced Aβ generation by PS2-γ-secretase.^43^ Lastly, the only evidence we are aware of, where endogenous PS expression has been compared utilised radioactive methionine labelling to correlate PS1 and PS2 antibody signals, and showed that in murine blastocyte derived membranes and cells, PS1-γ-secretase generated more Aβ than PS2-γ-secretase.^44^

Given the observed differences in tissues/cells, it is crucial to resolve the limitations of directly comparing cellular expression of PS1 and PS2 protein units and understand how this relates to γ-secretase activity. In this study, we investigated the activity of PS1 and PS2 in relation to the expression levels of these proteins, with an overarching hypothesis that PS1 does not have greater activity than PS2. We address this hypothesis using two approaches 1) Myc-tagging of the PS N-terminus to allow for detection of exogenous PS1 and PS2 via the same antibody, and 2) development of a novel PS1/2 fusion standard to enable for the first-time absolute quantitative assessment of endogenous PS1 and PS2 protein using specific antibodies. Our results demonstrate that in both the exogenous and endogenous PS expression systems, PS1 and PS2 are not equally expressed, and when PS expression is accounted for, PS2 is at least as active as PS1 at processing APP and Notch in HEK cells.

## MATERIALS AND METHODS

### Mammalian cell culture

All cell lines were cultured in Dulbecco’s Modified Eagle Medium (Sigma D5671) supplemented with 1 mM Sodium Pyruvate (Sigma S8636), 1 mM L-Glutamine (Sigma G7513), 100 units/ml Penicillin 0.1 mg/ml Streptomycin (Sigma P4333), and 10% v/v Foetal Bovine Serum (Serana FBS-Au-015). Cells were incubated at 37°C with 5% v/v atmospheric carbon dioxide.

### CRISPR Presenilin knockout in HEK-293

To generate HEK PS2+ cells, PS1 was knocked out of HEK 293 cells using Presenilin 1 CRISPR/Cas9 KO plasmid in conjunction with Presenilin 1 HDR Plasmid from Santa Cruz Biotechnology (sc-401227 and sc-401227-HDR) as per supplier protocol. Briefly HEK 293 cells were plated in 6-well plates (1.0 x 10^6^ cells/well), 24 hours prior to transfection. When cells were at approximately 80% confluency 1.25 µg each of the CRISPR/Cas9 KO and HDR plasmids were transfected using Lipofectamine 3000 (Invitrogen L3000015) and cells transfected as per manufacturer instructions. Cells were then incubated overnight after which the media was changed. At 48 hrs post transfection cells were sorted for GFP positive and RFP positive cells and cultured in puromycin selection media (0.25 µg/ml). Cells were selected for 8 days with media replacement every 48hrs.

PS2 knockout was completed using the pSp-Cas9-(BB)-2A-GFP vector and methodology previously described by Ran et al,^45^. Guide RNA sequences were designed using ChopChop.com.au, two guide sets were used in combination to generate the PS2KO in the cells. The guide sequences used were ^5’^GCTCCCCTACGACCCGGAGA^3^’ and ^5’^ACGATCATGCACAGAGTGAC^3’^. 10 µM each of sense and antisense synthesised oligonucleotides with appropriate flanking sequences,^45^ were phosphorylated using T4 PNK (NEB M0201) in a 10 µl reaction as per manufacturer protocol. The reaction was incubated at 37°C for 30 min, before heating to 95°C for 5 min followed by a temperature ramp of 1°C per min until reaching 25°C to anneal the oligonucleotides. Oligonucleotides were ligated into pSp-Cas9-(BB)-2A-GFP plasmid that had been linearised by digestion with BbsI-HF (NEB R3539) and gel purified (Bioline BIO-52060) as per supplier protocols. Briefly 2 µM of dsDNA guide was ligated into 10 ng of pSp-Cas9-(BB)-2A-GFP using 400 units T4 DNA ligase (NEB M0202) and incubated for 16 hrs at 16°C. 5 µl of ligation product was transformed into chemically competent E.coli XL10 cells and grown overnight on agar plates supplemented with 100 µg/ml ampicillin. Individual colonies were selected and cultured overnight in 5ml Luria Broth supplemented with 100 µg/ml ampicillin, plasmids were extracted (Bioline BIO-52057) and correct insertion of the guide RNA sequence confirmed by sequencing.

To generate HEK 293 PS1+ and HEK 293 PSnull cells, we seeded HEK 293 PS1+PS2+ and HEK 293 PS2+ cells in 6-well plates at 1.0x10^6^ cells per well 24 hours prior to transfection. 1.25 µg each of the two guide plasmids were prepared using Lipofectamine 3000 (Invitrogen L3000015) and cells transfected as per manufacturer instructions. Cells were incubated for 24 hours, after which they were sorted using a BD FACSJazz cell sorter at 1 cell per well into 96 well plates, gated for medium GFP intensity and no/minimal Propidium Iodide intensity. Monoclonal populations were expanded and screened for PS1 and PS2 protein expression, and selected clones were further screened for substrate processing (Fig S1). One representative clone was selected for subsequent experiments.

### Plasmid construct generation

All plasmid constructs used for transient transfection of PS and substrate proteins were generated in the backbone vector pIRES2-AcGFP1 (Takarabio). Human PS1 and PS2 cDNA sequence with Myc N-terminal tags, human APP695Swe and human Notch1 (lacking the extracellular domain (21-1713bp) termed ΔEhNotch1) sequences were cloned into pIRES2-AcGFP1 vector linearised via digestion by restriction enzymes at sites EcoRI and BamHI (PS1, PS2 and ΔEhNotch1) and SalI and XmaI (APP695Swe). After sequence confirmation 50 ml cultures were grown in Luria Broth and plasmids extracted (BioRad 7326120).

### Transient transfection

Transient transfection experiments were performed in 6-well plates, with cells seeded at 4.0 x 10^5^ per well. Prior to plating cells, plates were coated overnight with 50 µg/ml Poly-L-Lysine (Sigma P9155) to improve adherence of PS knockout cell lines. 24 hours after plating the media was replaced with antibiotic free media and transfected using Lipofectamine 3000 (Thermo Fisher L3000015) (for co-transfection of PSnull cells (Fig 2 & 3) or PEI Max (3 µg PEI per 1 µg DNA) (Polysciences 24765) (for substrate only transfection for investigating endogenous PS (Fig 5 & 6). Where substrate (pIRES2-AcGFP1-APP695Swe or pIRES2-AcGFP1-ΔEhNotch1) and presenilin (pIRES2-AcGFP1-Myc-PS1 or pIRES2-AcGFP1-Myc-PS2) co-transfection was undertaken in PSnull cells, the vectors were used in a PS:Substrate per unit ratio of 1:3, such that the total amount of DNA transfected was 500ng. For transfection of substrate only to investigate endogenous PS activity, the same amount of substrate vector was transfected as per co-transfection assays. Cells were incubated for 24 hours after which conditioned media, from the APP695Swe transfections, were collected for subsequent analysis by ELISA and the whole cell lysates harvested. Briefly, media was collected into microfuge tubes, centrifuged at 17,000 g for 5 min, the supernatant transferred into a clean tube and snap frozen in liquid nitrogen and stored at -80°C. For lysate collection, media was aspirated, and plates washed twice with cold PBS and aspirated. Cells were scrapped into 100 µl of RIPA lysis buffer (Astral Scientific 786-490) supplemented with protease inhibitor cocktail (Roche 11697498001) and transferred to microfuge tubes. Lysate samples were incubated for 1 hr at 4°C with rotation, centrifuged at 14,000 g for 10 min at 4°C, and the supernatant collected and stored at -20°C.

### Quantitative PCR

Cells were grown to confluency in 6-well plates and harvested for mRNA extraction. Briefly, plates were washed with cold PBS and cells scrapped and collected. Cells were centrifuged at 100g for 5min, after which supernatant was aspirated. RNA was extracted using ISOLATE II RNA Mini kit as per manufacturer instructions (Bioline BIO-52072) and RNA concentration and quality determined by Nanodrop (Thermo Fisher). 0.5 µg of RNA was used to generate cDNA using Tetro cDNA synthesis kit (Bioline BIO-65043) in a 10 µl reaction using a 1:1 ratio of random hexamer and oligo (dT)18 primer mix as per manufacturer instructions. Resultant cDNA samples were diluted to a final volume of 100 µl for use in qPCR. GoTaq qPCR master mix (Promega A6001) was used in a final reaction volume of 20 µl. For all genes, diluted cDNA solution (2 µl) was used in 20 µl reactions, primer details are listed in Table 1, and were designed using NCBI Primer-BLAST,^46^ except GAPDH primers.^47^ Each biological replicate was run in technical triplicate using the Applied Biosystems Viia7 real-time PCR system, and average Ct values determined for each gene, hUBC and hGAPDH reference genes were used for normalisation. Gene expression levels were calculated using the Pfaffl method,^48^ and expression relative to PS1+PS2+ cells determined.

**Table 1:**
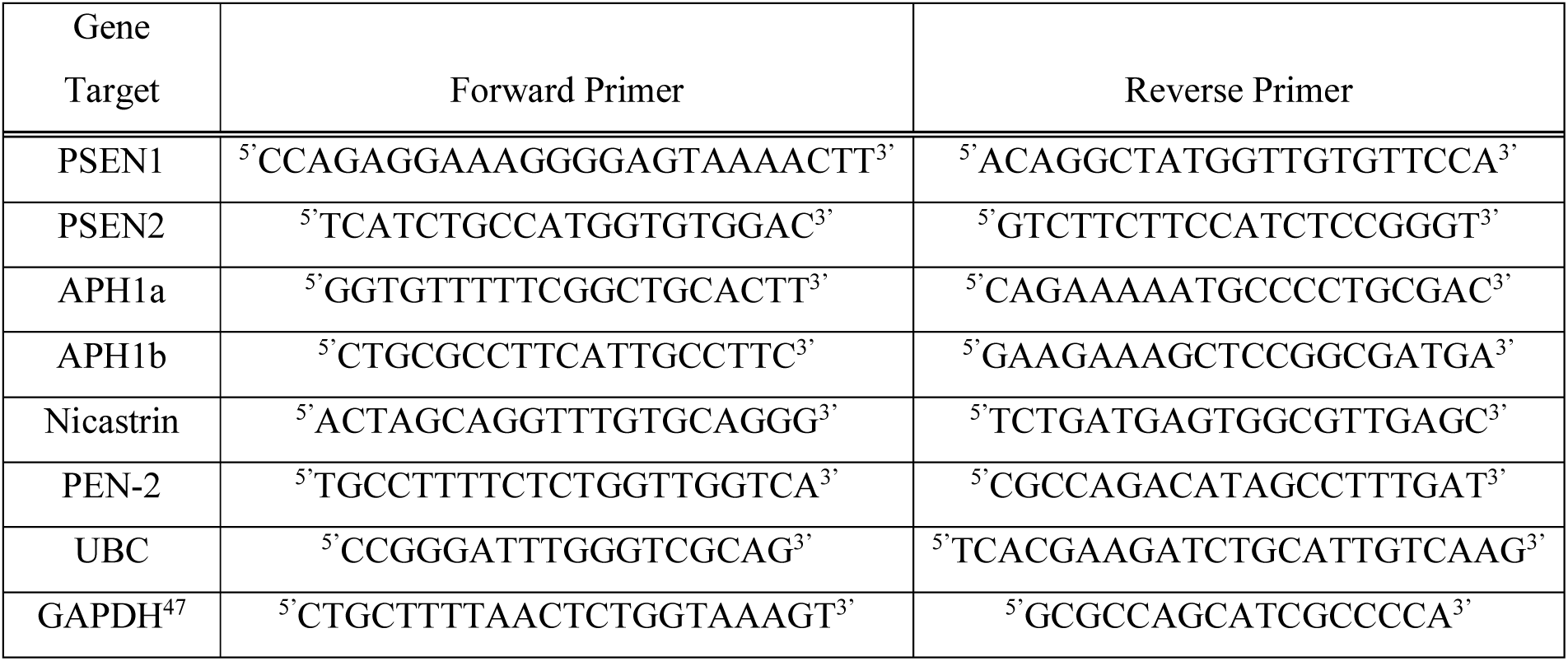
qPCR primer sequences

### Immunoblotting

Total protein concentration of cell lysates was determined using micro-BCA kit (Thermo Fisher 23235). Presenilin and APP proteins were separated with 12% v/v acrylamide, Tris-Tricine gel chemistry, and Notch, Pen-2, Aph1a, Nct proteins were separated with 8-10% v/v acrylamide, Bis-Tris gel chemistry (Invitrogen Surecast system). Samples were prepared using either 4x Tris-Tricine sample buffer (16% w/v SDS, 200 mM tris, 48% v/v glycerol, 0.5% w/v Coomassie G-250) or 4x LDS sample buffer (Thermo Fisher B0007) as appropriate, reducing agent and heat treatment conditions vary dependent on the protein of interest (see Table 2). Samples were vortexed for 30 seconds, heated treated as appropriate, centrifuged at 17,000 g for 5 minutes, then electrophoresed at 100v for 1 hour 45 minutes. Proteins were transferred to 0.2 µm nitrocellulose membrane (BioRad 1620112), via wet transfer method using Tris-Gylcine buffer (19.2 mM glycine, 2.5 mM tris, 20% v/v methanol) at 150 mA for 16 hours at 4°C. Membranes were stained with ponceau S (1% w/v Ponceau S, 5% v/v acetic acid) for 5 minutes to assess transfer quality before destaining with boiled TBS (2 mM tris, 1.5 mM NaCl). Membranes were subsequently incubated in blocking buffer as appropriate for the primary antibody used for 1 hour at room temperature with agitation. Membranes were incubated in primary antibody (All antibody conditions and details available in Table 2) overnight at 4 °C. Membranes were subsequently washed three times in TBS-tween (0.05%) for 10 minutes with agitation. Subsequently the membranes were incubated with appropriate secondary antibody, anti-mouse or anti-rabbit IgG HRP conjugated secondary antibody (Thermo Fisher 31430, 31460), diluted at 1:20,000 in 0.5% w/v non-fat dry milk, for 1 hour at room temperature with agitation. Membranes were then washed again three times in TBS-tween, followed by a 5-minute wash with TBS. Membranes were incubated in either Clarity ECL (BioRad 1705061) or Prime ECL (Cytiva GERPN2232) (see Table S2) for 5 minutes as per manufacturer instructions, then imaged on a BioRad ChemiDoc MP system.

**Table 2:**
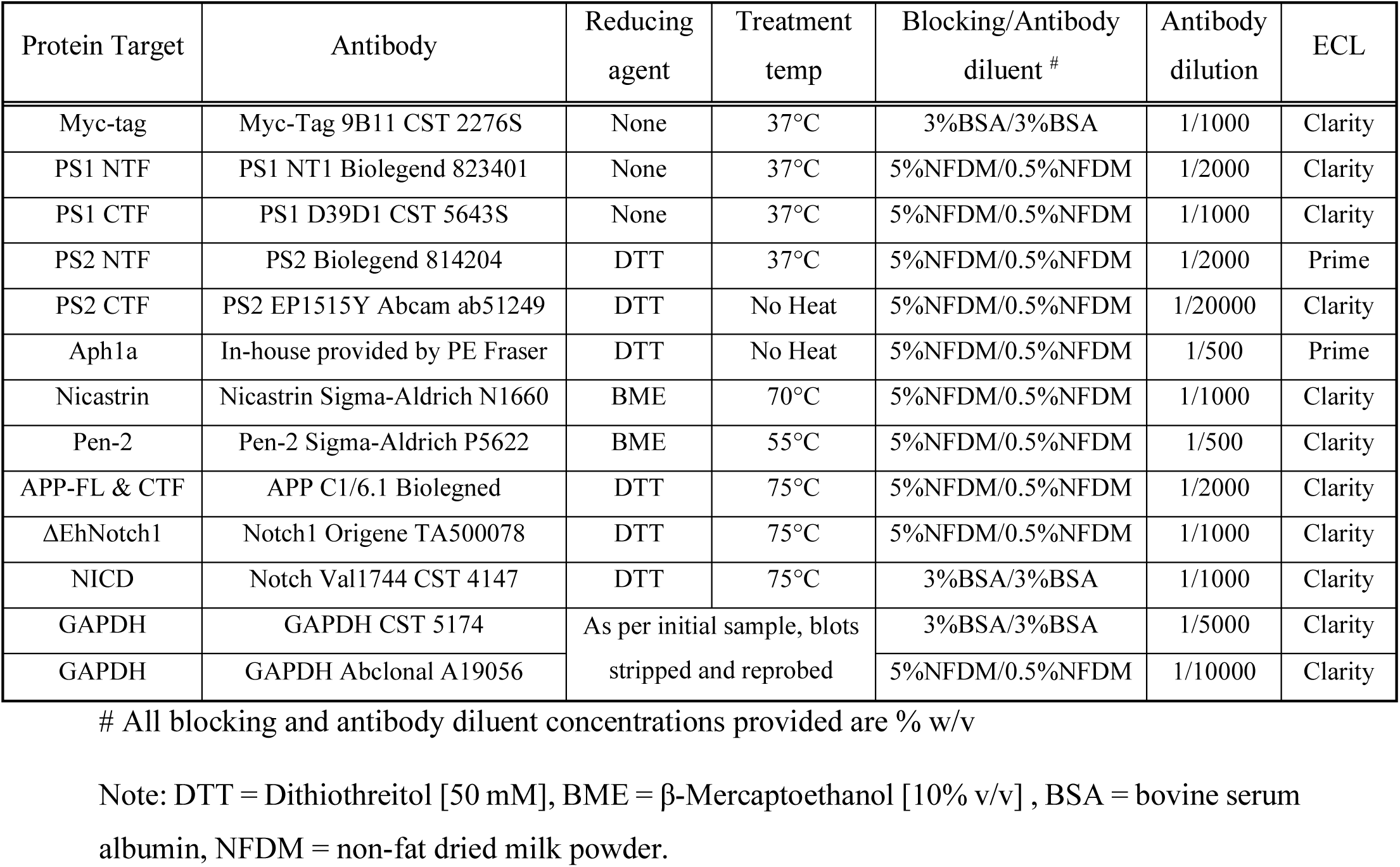
Antibody conditions for immunoblotting

### PS1/2 fusion standard and absolute PS1 and PS2 quantitation

The presenilin fusion standard (PS-Std) (see results section, Fig 4A) was recombinantly generated in E. coli and purified by GenScript. The final size of the protein including tags is 30.7 kDa, which equates to 5.10x10^-11^ ng per protein unit. In order to use the standard to quantify the number of endogenous PS protein units a 5 to 6-point standard curve was generated on every PS immunoblot, the actual range used was determined empirically and is dependent on the total protein of sample used, the protein fragment being detected i.e., PS1 or PS2, NTF or CTF, and the specific antibody used. The ng of PS-Std is converted to protein units of PS-Std as follows; protein units PS-Std = ((ng PS-STD) x 90%)/5.10x10^-11^ (note the PS-Std purity was determined to be 90% in quality control report from supplier). This value is then plotted against the corresponding band intensity densitometry units quantitated from the immunoblot, to generate a standard curve, using multiple replicates. The standard curve line of best fits equation was used to determine the PS1 or PS2 protein units in the sample. Therefore, the total PS protein units in PS1+PS2+ cells will be the sum of PS1 and PS2 protein units.

### ELISA

Invitrogen ELISA kits were used to detect Aβ40 (KHB3482) and Aβ42 (KHB3442) in conditioned media as per manufacturer instructions. For detection of Aβ40 from conditioned media endogenous PS activity samples were diluted 1/6, while exogenous PS activity samples were diluted 1/3. No dilution was necessary for detection of Aβ42 in conditioned media.

### Statistics

All statistical analysis was completed using GraphPad Prism 9.5.0. Four to six experimental replicates were completed for all assays. Statistical significance was determined via unpaired T-test where only two groups were examined, where more than two groups are examined one-way ANOVA or two-way ANOVA analysis with Holm-Šidák’s multiple comparison tests are used as appropriate.

## RESULTS

### Presenilin knock-out cell line generation and characterisation

To evaluate and compare the influence of exogenous and endogenous PS1 or PS2 on γ-secretase, presenilin knockout cell lines derived from HEK-293WT (PS1+PS2+) were generated. The cell lines included those lacking PS1 but retaining PS2 expression, lacking PS2 but retaining PS1 expression, or lacking both PS1 and PS2 protein expression. These cell lines were referred to as PS2+, PS1+, and PSnull, respectively. Monoclonal populations were assessed for PS1 and PS2 expression as well as γ-secretase processing of APP and Notch1 substrates (Fig S1), and the clone representative of average substrate processing selected for use in subsequent assays.

The expression of the components of γ-secretase is regulated by the formation of the enzyme complex.^1, 49–51^ Therefore, to assess the effect of the absence or presence of PS on the expression of γ-secretase components and to further characterise the cell lines, expression of PSEN1, PS2, Aph1, Pen-2 and Nct was assessed at transcript (Fig 1A) and protein levels (Fig 1B-F). As expected, compared to the wild-type, *PSEN1* and *PSEN2* mRNA expression is absent or markedly reduced in cells where the specific PS has been ablated. In the single PS knockout cell lines, an increase in mRNA expression was observed for the alternate homologue, indicating that loss of one PS homologue is causing a compensatory increase in transcript expression of the alternate homologue. For all other components of γ-secretase, mRNA expression is significantly reduced in the PS2+ and PSnull cell lines, while no significant changes were observed in the PS1+ cell line.

**Figure 1.**
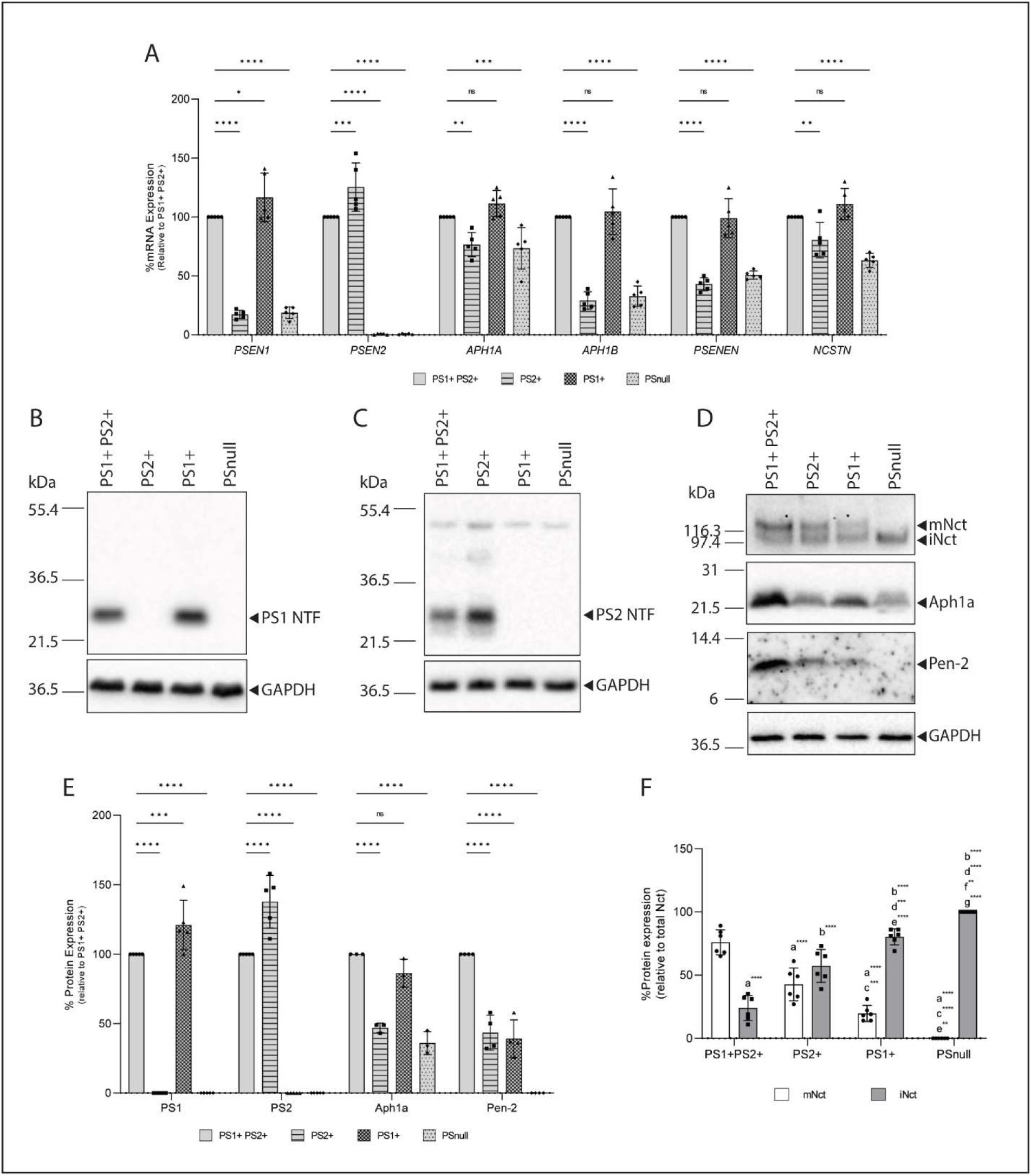
Characterisation of HEK 293 PS1+PS2+, PS1+, PS2+ and PSnull cell lines. Cell lines were generated by CRISPR-Cas9 knockout of PSEN1 and/or PSEN2 from HEK 293 cell lines. Representative clone of each cell line selected for further analysis by mRNA expression of γ-secretase subunits PS1, PS2, Aph1a, Aph1b, Pen-2 and Nct and presented relative to PS1+PS2+ cell line expression (A). Whole cell lysates were analysed by immunoblot for detection of PS1 protein (B) PS2 protein (C) and Aph1a, Pen-2 and Nct proteins (D). Protein expression levels were quantitated by densitometry analysis and are presented relative to PS1+PS2+ cell line expression for PS1, PS2, Aph1a and Pen-2 (E). Both mature Nct (mNct) and immature Nct (iNct) were quantitated and the percentage of each relative to total Nct calculated (F). Values shown are mean ± SD of n = 3-6 independent experiments. Two-way ANOVA with Holm-Šidák’s multiple comparison was completed for (A, E, F). For (F) a = significantly different to mNct PS1+PS2+, b = significantly different to iNct PS1+PS2+, c = significantly different to mNct PS2+, d = significantly different to iNct PS2+, e = significantly different to mNct PS1+, f = significantly different to iNct PS1+, g = significantly different to mNct PSnull. For all quantitated data; ns = P > 0.05, * = P < 0.05, ** = P < 0.01, *** = P < 0.001, **** = P <0.0001.

PS1 and PS2 protein expression (Fig 1B, C, E) is commensurate with observed mRNA expression. No PS1 or PS2 protein is detected in cells where the appropriate PS has been accordingly knocked out. Increases in PS1 and PS2 protein are observed in the PS1+ and PS2+ cell lines respectively, in line with the observed mRNA results. Aph1a protein levels similarly align with the mRNA results (Fig 1D, E). However, Pen2 protein levels differ from the mRNA expression profile, with significant reduction in protein levels evident in both the PS1+ and PS2+ cell lines, and no protein detected in PSnull cells (Fig 1D, E).

In all cell lines except for PSnull, nicastrin was detected as two protein bands, representing immature and mature protein (Fig 1D, F). In the PSnull cells, nicastrin was detected as immature protein only, as has been previously reported in double knockout cells^37, 42^ and consistent with the requirement of presenilin for maturation of nicastrin.^50^ Notably, mature vs immature Nct levels vary in a PS dependent manner, consistent with previous reports.^32, 37^ Where PS1+PS2+ cells have approximately 4-times more mature Nct, PS2+ cells have approximately equal levels of mature and immature Nct, and PS1+ cells have approximately 4-times more immature Nct.

### Exogenous PS expression highlights difference in PS levels and subsequently higher PS2 activity

We proceeded to investigate the processing of hAPP695Swe and hNotch1 in PSnull cells with the intent of directly comparing exogenously expressed PS1 and PS2 for the assessment of γ-secretase activity in the absence of endogenous presenilin. This was achieved by using PS1 and PS2 constructs N-terminally tagged with Myc. N-terminal tagging has been previously used for exogenous PS expression,^32, 41^ while C-terminal tagging would be unsuitable as this region interacts with the Aph1 component of γ-secretase.^52^ Exogenous, myc-tagged, PS1 (exPS1) or PS2 (exPS2) was co-transfected with either hAPP695Swe or ΔEhNotch1 at a ratio of 1:3 (PS:Substrate). The amount of exPS used in the transfections was titrated to reduce the amount of unincorporated full-length protein, while retaining maximum PS-NTF levels (Fig S2).

APP processing was assessed via immunoblotting (Fig 2A) of whole cell lysates from co-transfection with either PS1 or PS2 and APP695Swe. APP695 full length and APP-CTF levels were quantitated and expressed as the ratio of APP-CTF/APP as an initial indicator of γ-secretase activity. APP-CTF protein accumulation was significantly reduced with the co-expression of either exPS1 or exPS2 compared to control; notably, APP-CTF accumulation was 2.0-fold lower with exPS2 compared to exPS1 (Fig 2B). In contrast, the Aβ40 and Aβ42 levels in conditioned media detected by ELISA, were significantly higher when APP695Swe was co-expressed with exPS1 (Fig 2C, D).

**Figure 2.**
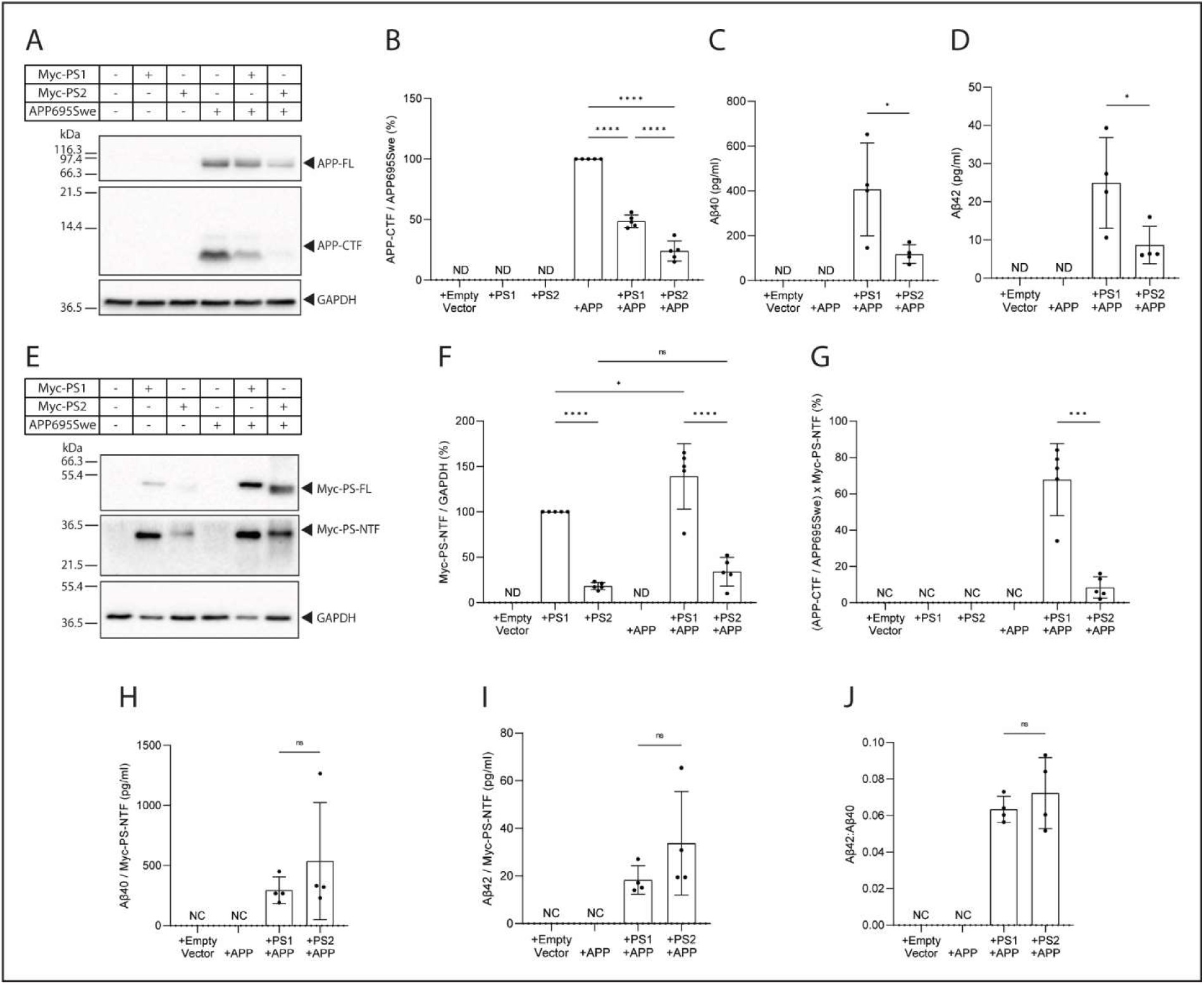
APP processing by exogenous myc-tagged PS1 and PS2 in PSnull cells. Both APP695Swe and myc-tagged PS were transiently co-expressed in PSnull cells to assess APP processing and directly compare PS1 and PS2 expression to enable the effect of variable expression to be considered. Whole cell lysates were assessed via immunoblot to determine APP695Swe and APP-CTF protein levels (A) and the accumulation of APP-CTF/APP695Swe quantitatively determined by densitometry assessment and presented relative to PSnull cells transfected with APP695Swe only (B). Conditioned media was collected concurrently with whole cell lysates for analysis of Aβ40 (C) and Aβ42 (D) levels (pg/ml) by ELISA. Exogenous PS1 and PS2 expression was directly compared by immunoblotting using antibody directed against the myc-tagged N-terminus of exPS1 and exPS2 (E). Note that to enable simultaneous detection of exPS1 and exPS2 the total protein loaded was adjusted such that for exPS1 transfected lysates 10 µg total protein was loaded, while for exPS2 transfected lysates 30 µg of protein was loaded. Myc-PS-NTF levels were quantitated by densitometry analysis and normalised for GAPDH to account for the different amounts of total protein loaded between exPS1 and exPS2 samples (F). APP-CTF, Aβ40 and Aβ42 were subsequently normalised for Myc-PS-NTF levels to account for variable PS1 and PS2 expression (G-I). Aβ42:Aβ40 ratio was calculated (H). Values shown are mean ± SD of n = 4-5 independent experiments. Statistical tests applied were unpaired t-test for (C, D, G-J) and ordinary one-way ANOVA with Holm-Šidák’s multiple comparison for (B, F) where ns = P > 0.05, * = P < 0.05, ** = P < 0.01, *** = P < 0.001, **** = P <0.0001.

When investigating exogenous PS expression, it was initially noted that Myc-PS2 NTF expression levels were dramatically lower than Myc-PS1 NTF expression; indeed, the total protein loaded had to be adjusted in order to detect both proteins on the immunoblot (Fig 2E). Normalised quantitation of the Myc-PS-NTF showed that in the absence of APP695Swe expression, exPS1 expression is 5.5 fold higher than exPS2. In the presence of APP695Swe, exPS1 expression significantly increases, and a trend towards an increase is observed with exPS2 expression. (Fig 2F). Having quantitated exPS1-NTF and exPS2-NTF expression levels using myc-tag detection, enabling direct comparison, we were able to use this data to normalise APP695Swe substrate processing activity to determine the specific contributions of exPS1- and exPS2-γ-secretase. Consequently, we determined that exPS1-γ-secretase accumulates 8.1-fold more APP-CTF than exPS2-γ-secretase (Fig 2G). Similarly, Aβ40 and Aβ42 levels were normalised to exPS-NTF expression, demonstrating no significant difference in Aβ40 or Aβ42 levels (Fig 2H, I). Additionally, no significant difference was observed in the Aβ42:Aβ40 ratios between exPS1 and exPS2 co-expression systems (Fig 2J).

To determine the effect of Presenilin expression on Notch1 cleavage, exPS1 or exPS2 was co-transfected with ΔEhNotch1. To assess substrate processing, the Notch1 ICD (NICD), which directly reflects γ-secretase activity, and ΔEhNotch1 proteins were detected via immunoblotting. The NICD/ΔEhNotch1 ratio was assessed, initially without consideration of exPS-NTF expression. As expected, no NICD product was detected in the absence of PS expression, and exPS1 co-transfection led to a 2.7-fold higher NICD/ΔEhNotch1 ratio than exPS2 co-transfection (Fig 3A,B), indicating more NICD product generated. Like that done for APP cleavage, myc-PS-NTF levels in the absence or presence of ΔEhNotch1 expression was quantitated and used to normalise the activity data (Fig 3). In the absence of substrate overexpression, 4.5-fold more exPS1-NTF is expressed than exPS2-NTF (Fig 3 C and D). In the presence of substrate, co-expression of exPS1 leads to increased PS1-NTF levels, while no change is observed in PS2-NTF when exPS2 is expressed (Fig 3 C and D). These results indicate an up-regulation of PS1 but not PS2 when Notch substrate is expressed. Using the myc-PS expression data to normalise the NICD/ΔEhNotch1 ratio revealed that, despite the up-regulation in PS1-NTF, the ratio (indicative of more NICD production) is increased in cells expressing exPS2 only, compared to those cells expressing exPS1 only (Fig 3E).

**Figure 3.**
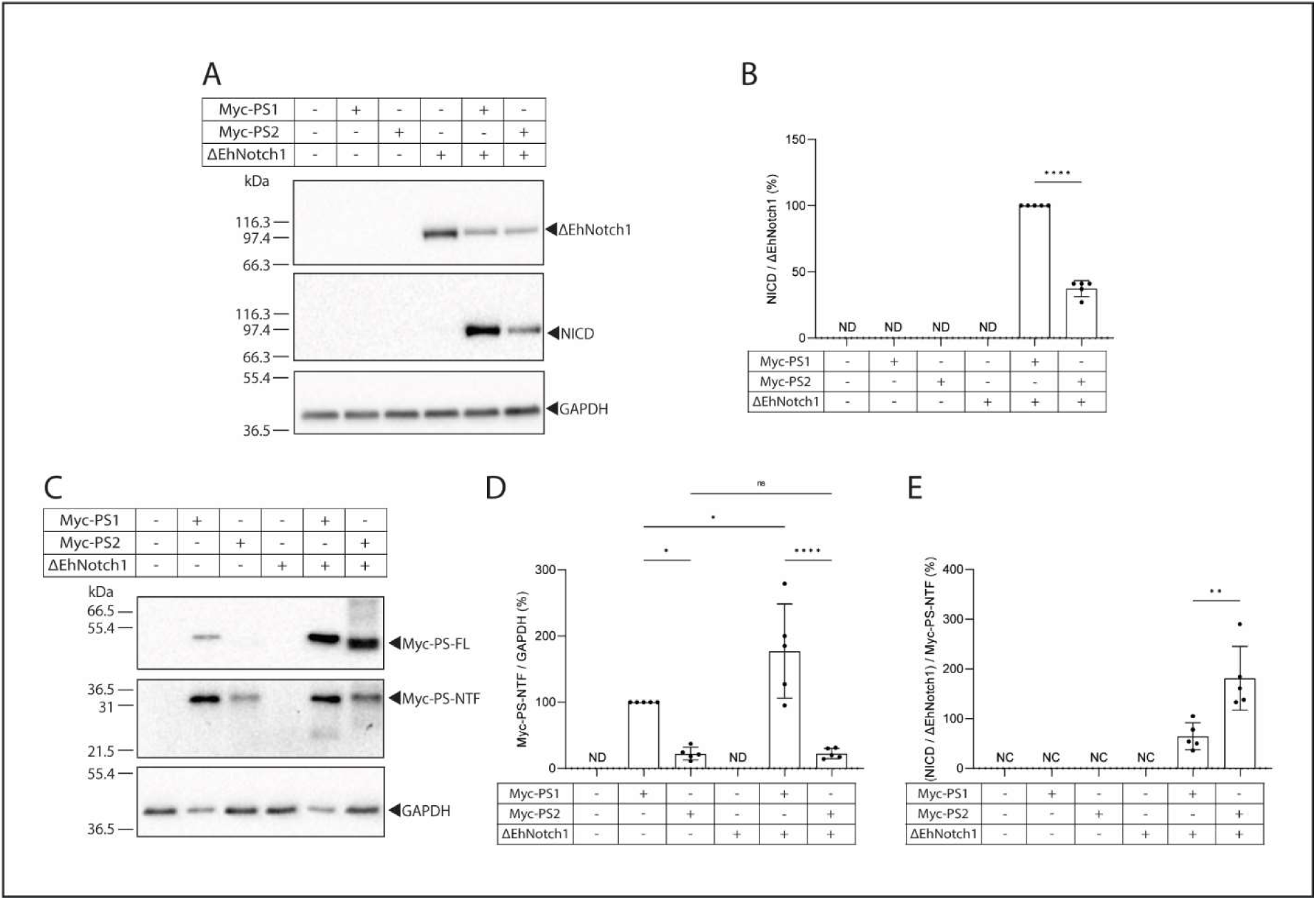
Notch1 processing by exogenous myc-tagged PS1 and PS2 in PSnull cells. Both ΔEhNotch1 and myc-tagged PS were transiently co-expressed in PSnull cells to assess APP processing and directly compare PS1 and PS2 expression to enable the effect of variable expression to be considered. Whole cell lysates were assessed via immunoblot to determine ΔEhNotch1 and NICD protein levels (A) and the level of NICD/ΔEhNotch1 quantitatively determined by densitometry assessment and presented relative to exPS1 transfection with ΔEhNotch1 (B). Exogenous PS1 and PS2 expression was directly compared by immunoblotting using an antibody directed against the myc-tagged N-terminus of exPS1 and exPS2 (C). Note that to enable simultaneous detection of exPS1 and exPS2 the total protein loaded was adjusted such that for exPS1 transfected lysates 10 µg total protein was loaded, while for exPS2 transfected lysates 30 µg of protein was loaded. Myc-PS-NTF levels were quantitated by densitometry analysis and normalised for GAPDH to account for the different amounts of total protein loaded between exPS1 and exPS2 samples (D). NICD was subsequently normalised for Myc-PS-NTF levels to account for variable PS1 and PS2 expression (E). Statistical tests applied were unpaired t-test for (B, E) and ordinary one-way ANOVA with Holm-Šidák’s multiple comparison for (D) where ns = P > 0.05, * = P < 0.05, ** = P < 0.01, *** = P < 0.001, **** = P <0.0001.

Overall, the findings indicate that exogenous expression of PS1 and PS2 do not lead to comparable expression of the γ-secretase incorporated myc-PS-NTF proteins. Notably, when the difference in expression is accounted for, exogenously expressed PS2 is more active than PS1 at cleaving APP695Swe and ΔEhNotch1.

### PS1/2 fusion standard: A method for absolute quantitation of endogenous PS1 and PS2

Recognising that significantly more PS1 is expressed than PS2 in an exogenous system, we sought to understand whether this PS expression profile (and effects on γ-secretase activity) was an artifact or recapitulated the endogenous expression profile. The ability to quantitatively compare endogenous PS1 (enPS1) and PS2 (enPS2) expression levels remains a challenge, as PS1 and PS2 are detected by different antibodies, with no commercially available antibody able to detect both homologues. To facilitate this, we designed a presenilin fusion standard (PS-Std). The PS-Std incorporates residues from the N-terminal sequence and the cytoplasmic loop of human PS1 and PS2. These regions are hydrophilic, non-transmembrane regions that contain the epitopes for several commercially available antibodies (Fig. 4A, Table S3).

PS1 and PS2 antibodies that detect either the N-terminal fragment (NTF) or C-terminal fragment (CTF) were used to probe for the PS-Std via immunoblot (Fig 4). All antibodies detect a single band on a tris-tricine gel under denatured conditions across the mass range of PS-Std used. The theoretical size of the PS-Std is 30.7kDa, however, the standard migrates at approximately 37 kDa, likely due to it being relatively acidic (pI=4.15).^53, 54^ When probed with unrelated antibodies, anti-GAPDH and anti-GSK3β, no bands were detected in the PS-std samples (Fig S3), confirming that the PS-std contains PS epitopes specific to PS1 and PS2 antibodies. Having validated antibody detection of the standard, we set out to quantitate endogenous PS expression levels in the HEK presenilin knockout cell lines generated.

To quantitatively assess expression, varying amounts of the PS-Std underwent SDS-PAGE along with whole cell lysate samples from the cell lines and subsequently detected via immunoblotting using PS1-NTF, PS1-CTF, PS2-NTF and PS2-CTF antibodies (Fig 4B-E). The densitometry results for the PS-Std were used to generate standard curves for each antibody and set of immunoblot replicates. Due to protein size differences between the PS-Std, and the NTF and CTF of PS1 and PS2, we did not simply determine the equivalent mass of standard, but rather, determined the number of PS protein units. This was achieved by calculating the number of PS-Std units of protein per ng of PS-Std (given one PS-Std unit is 5.10x10^-11^ ng) and plotting against the corresponding densitometry values (Fig S4). The equations from the resultant standard curves were used to convert the densitometry results to the number of PS1 or PS2 protein units. This value was subsequently normalised for total protein loaded on the PAGE, to determine the PS protein units per µg total protein.

Importantly, no significant differences were observed in any of the cell lines between the expression levels of the NTF and CTF proteins for either PS1 or PS2 (Fig 4F). This result further supports the use of the PS-Std for quantitation, as γ-secretase is known to contain components in a stoichiometric ratio of 1:1:1:1,^55^ and PS NTF and CTF fragments are tightly regulated at a 1:1 ratio.^56^ We calculated the difference in PS1 and PS2 expression, and determined that in the wild-type (PS1+PS2+ cells), PS1 expression is 5.2-fold higher than PS2 expression, which closely aligns with the exogenous expression profile determined above (see results section *Exogenous PS expression highlights difference in PS levels and subsequently higher PS2 activity*), reflecting that PS expression is tightly regulated at a homologue specific level.

**Figure 4.**
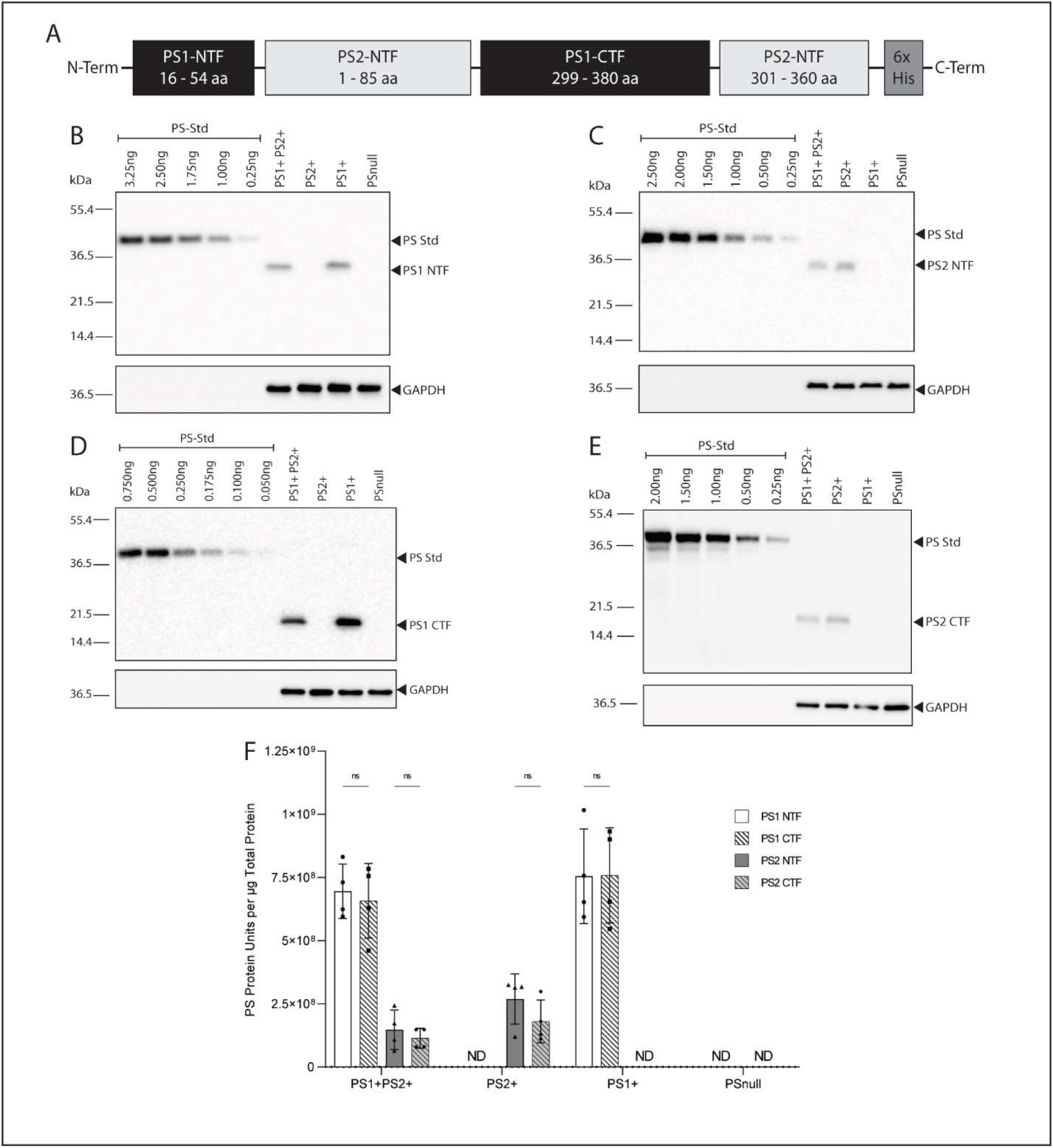
Validation of novel method to directly compare endogenous PS1 and PS2 expression. No commercially available antibody detects both PS1 and PS2, so to enable direct quantitation of endogenous PS1 and PS2 expression we developed a synthetic PS1/2 fusion protein (PS-Std) containing multiple epitope regions for several commercially available antibodies for use as a standard to enable comparative quantitation. The PS-Std contains protein sequences for the N-terminus and the cytoplasmic loop regions of PS1 and PS2 as shown schematically (A). A range of amounts of the PS-Std were immunoblotted alongside whole cell lysates from the PS1+PS2+, PS1+, PS2+ and PSnull cells to generate a standard curve using the same experimental conditions and probed with antibodies directed against the PS1 NTF (B), PS2 NTF (C), PS1 CTF (D) and PS2 CTF (E). Immunoblot bands underwent densitometry assessment for quantitation and the PS-Std densitometry results were used to generate a standard curve for each replicate set of immunoblots relative to the number of PS-Std protein units ((ng PS-STD) x 90%/5.10x10^-11^) (Fig S4). The resultant standard curve was used to calculate the amount of PS protein units detected in the whole cell lysate samples for each cell line (F). Values shown are mean ± SD of n = 4 independent experiments and analysed using two-way ANOVA with Holm-Šidák’s multiple comparison where ns = P > 0.05.

### Endogenous PS2 demonstrates greater activity in cleaving APP and equivalent activity in cleaving Notch

Having validated the method for quantitating endogenous PS protein levels, we next sought to investigate γ-secretase substrate processing by endogenous PS1- and PS2-γ-secretase in the HEK presenilin knockout cell lines. hAPP695Swe was transiently transfected into the cell lines, and whole cell lysates were harvested for immunoblotting 24 hrs after transfection; additionally, conditioned media was collected for the determination of Aβ generation.

APP-FL and APP-CTF levels were detected via immunoblotting and quantitated (Fig 5A, B). Levels of overexpressed APP-FL were noticeably variable between the cell lines (particularly in the absence of PS1), despite equal levels of GAPDH loading control. Consequently, APP-CTF accumulation was represented as a ratio of APP-FL, to measure PS-specific γ-secretase activity. In the PSnull cells, considerable APP-CTF accumulated as a result of lack of γ-secretase activity, due to the absence of PS. Comparatively, less than 2% APP-CTF was detected in PS1+PS2+ cells. While the accumulation of APP-CTF protein in both the PS2+ and PS1+ cells is significantly less than the PSnull cells, significantly higher APP-CTF levels were detected in PS2+ cells compared with the PS1+PS2+ cells. Conditioned media collected from hAPP695Swe transfections was analysed via ELISA, Aβ40 levels are significantly higher in the PS1+ cells compared to both the PS1+PS2+ and PS2+ cells, whereas both PS1+ and PS2+ cells generate increased Aβ42 compared to PS1+PS2+ cells (Fig 5C, D). These results are indicative of PS2-γ-secretase preferentially initiating the Aβ42 generation pathway, consequently leading to a higher Aβ42:Aβ40 ratio (Fig 5 K).

**Figure 5.**
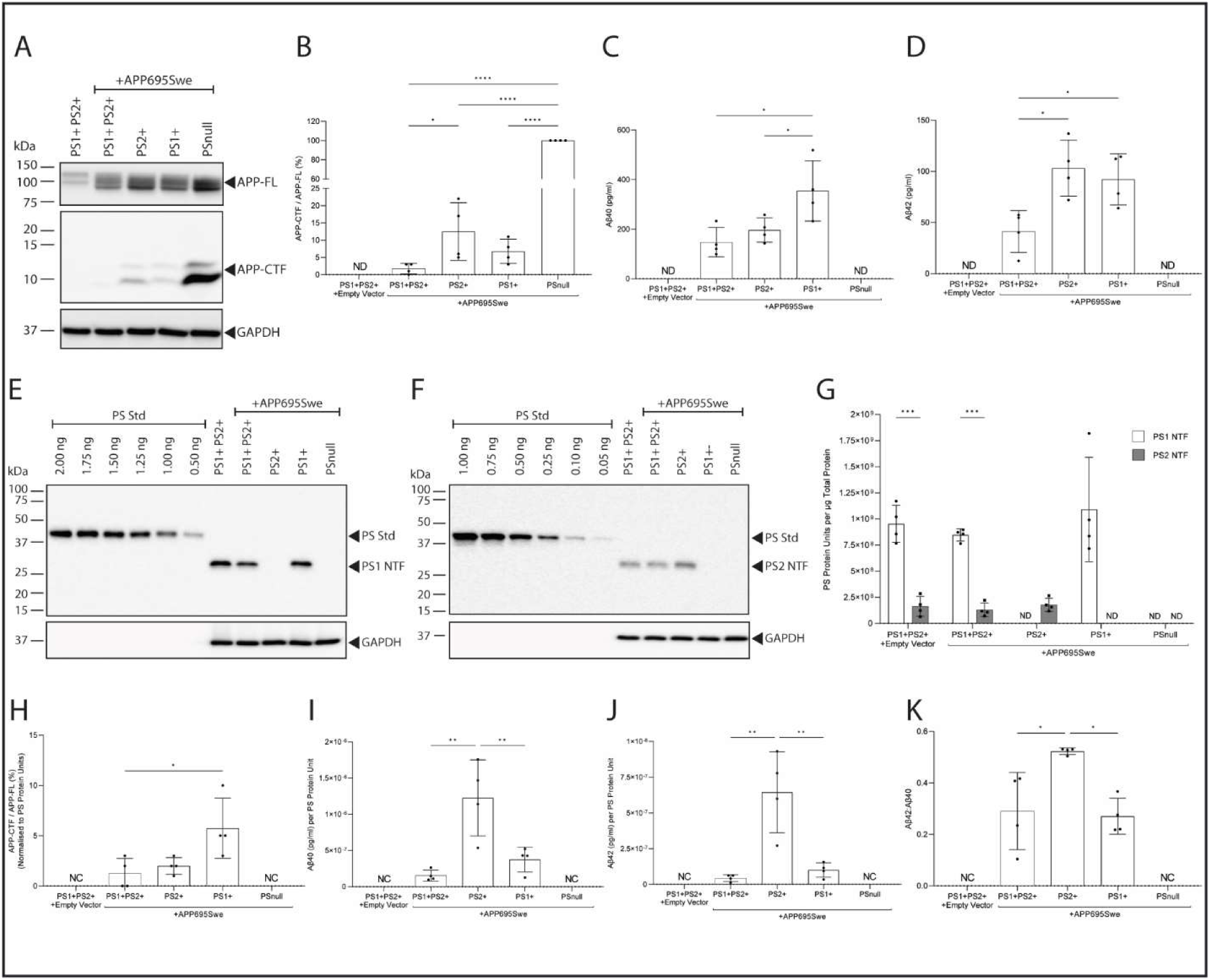
APP processing by endogenous PS1 and PS2 assessed in PS1+ and PS2+ cells. APP695Swe was transiently expressed in PS1+PS2+, PS1+, PS2+ and PSnull cells to assess APP processing by endogenous PS1 and PS2 and our novel PS-Std used to quantitatively determine PS1 and PS2 expression to enable the effect of variable expression to be considered. Whole cell lysates were assessed via immunoblot to determine APP695Swe and APP-CTF protein levels (A) and the accumulation of APP-CTF/APP695Swe quantitatively determined by densitometry assessment and presented relative to PSnull cells (B). Conditioned media was collected concurrently with whole cell lysates for analysis of Aβ40 (C) and Aβ42 (D) levels (pg/ml) by ELISA. Endogenous PS1 and PS2 expression was determined by immunoblotting using antibodies directed against the PS1 NTF and PS2 NTF (E, F). PS1 NTF and PS2 NTF levels were quantitated by densitometry analysis and the PS protein units determined using standard curves generated alongside the whole cell lysates (G). APP-CTF, Aβ40 and Aβ42 were subsequently normalised for PS protein units to account for variable PS1 and PS2 expression (H-J). Aβ42:Aβ40 ratio was calculated (K). Values shown are mean ± SD of n = 4 independent experiments. Statistical tests applied were ordinary one-way ANOVA with Holm-Šidák’s multiple comparison (B-D, H-K) and two-way ANOVA with Holm-Šidák’s multiple comparison for (G) where ns = P > 0.05, * = P < 0.05, ** = P < 0.01, *** = P < 0.001, **** = P <0.0001.

Endogenous PS1 and PS2 levels were detected via immunoblotting and quantitated against the PS-Std to determine PS protein units (Fig 5E-G). Expression levels of enPS1 was greater than enPS2 in PS1+PS2+ cells lacking (5.9-fold higher) or over-expressing hAPP695swe (6.5-fold higher) and no significant difference was observed in the level of either enPS1 or enPS2 in the absence of the alternate PS homologue. We next assessed substrate processing at a per PS unit level to allow for a more accurate assessment of γ-secretase activity. Prior to normalisation, it was observed that more APP-CTF accumulated in PS2+ cells, compared to PS1+ cells, which is indicative of less processing (Fig 5B). Normalising for PS units reveals that, when the lower levels of PS2 were considered, PS2+ cells show less APP-CTF compared to PS1+ cells, indicative of increased processing (Fig 5H). Taken together with the results of Aβ40 and Aβ42 (Fig 5I, J), these findings are indicative that PS2γ-secretase is more active at processing hAPP695swe (Fig 5I, J).

Furthermore, endogenous PS processing of Notch1 was investigated by transiently transfecting cell lines with the ΔEhNotch1 vector and collecting whole cell lysates after 24 hrs. NICD levels and ΔEhNotch1 levels were detected via immunoblotting, and the NICD/ΔEhNotch1 levels determined (Fig 6A,B). As expected, no NICD is detected in either the PS1+PS2+ cells transfected with the vector control or in the PSnull cells. Prior to any consideration of PS expression levels, there is no significant difference observed between the levels of NICD generated by PS1+PS2+ and PS1+ cells. The PS2+ cells, however, generate 3.0-fold less NICD than the PS1+PS2+ or PS1+ cells. On quantitating PS expression levels, significantly higher expression of enPS1 than enPS2 expression (4.8-fold higher enPS1) was observed in PS1+PS2+ cells (Fig 6C-E). No significant differences between enPS1 or enPS2 levels were observed in either PS1+ or PS2+ cells, compared with PS1+PS2+ cells. Subsequently, expressing NICD generation relative to PS protein units, no difference was observed between any cell lines (Fig 6F).

**Figure 6.**
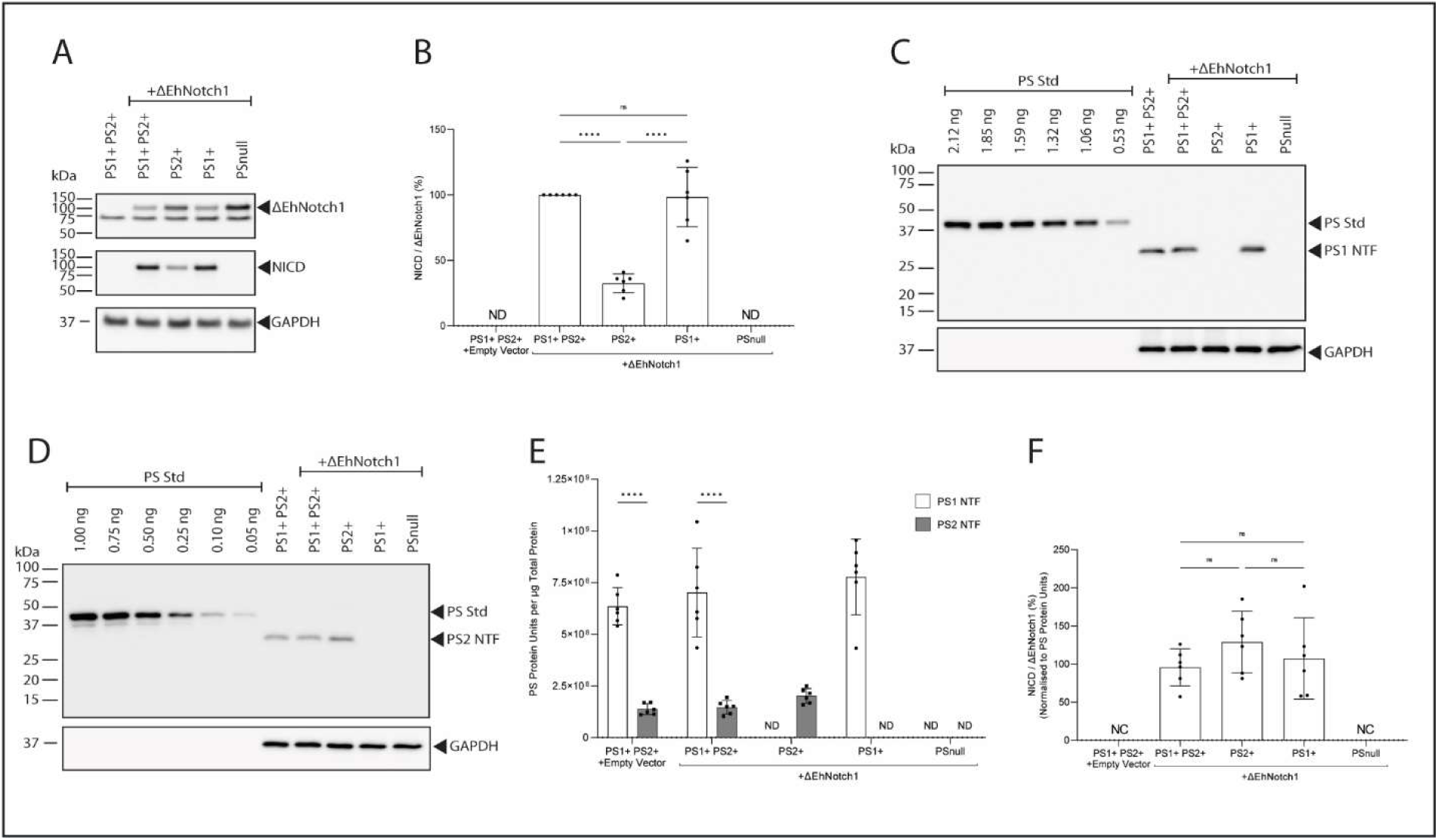
Notch1 processing by endogenous PS1 and PS2 assessed in PS1+ and PS2+ cells. ΔEhNotch1 was transiently expressed in PS1+PS2+, PS1+, PS2+ and PSnull cells to assess Notch1 processing by endogenous PS1 and PS2 and our novel PS-Std used to quantitatively determine PS1 and PS2 expression to enable the effect of variable expression to be considered. Whole cell lysates were assessed via immunoblot to determine ΔEhNotch1 and NICD protein levels (A) and the generation of NICD/ΔEhNotch1 quantitatively determined by densitometry assessment and presented relative to PS1+PS2+ cells (B). Endogenous PS1 and PS2 expression was determined by immunoblotting using antibodies directed against the PS1 NTF and PS2 NTF (C, D). PS1 NTF and PS2 NTF levels were quantitated by densitometry analysis and the PS protein units determined using standard curves generated alongside the whole cell lysates (E). NICD was subsequently normalised for PS protein units to account for variable PS1 and PS2 expression (F). Values shown are mean ± SD of n = 6 independent experiments. Statistical tests applied were ordinary one-way ANOVA with Holm-Šidák’s multiple comparison (B, F) and two-way ANOVA with Holm-Šidák’s multiple comparison for (E) where ns = P > 0.05, * = P < 0.05, ** = P < 0.01, *** = P < 0.001, **** = P <0.0001.

## DISCUSSION

The study of the specific contributions of PS1 and PS2 proteins to γ-secretase substrate processing, be it using endogenous PS or exogenous overexpression of PS, typically does not consider PS1 vs PS2 expression levels. Although the few studies that have directly compared presenilin expression show higher PS1 expression than PS2,^41, 44^ there is evidence that PS2 expression increases with age,^38, 40^ is associated with ADAD^28^ and increases in response to mutant PS1 expression,^36^ suggesting the importance of PS2 in AD pathogenesis. Our current study has developed and applied the use of new methods for the direct quantitation of both exogenous and endogenous PS expression and the findings highlight the importance of considering PS expression when interpreting γ-secretase activity. We show that in HEK 293 cells, there is approximately 5-times more PS1 than PS2 protein expression, likely as a result of the embryonic origin of this cell line.^38^ Importantly, we identify that this expression profile is retained when exogenous PS1 and PS2 are expressed after PS ablation. Finally, we have demonstrated that when PS expression is considered, PS2-γ-secretase processes at least equal amounts, if not more, APP and Notch1 than PS1-γ-secretase, dependent on the experimental system in use.

Our novel method for quantitating endogenous PS demonstrates that in HEK 293 cells, PS1 expression is significantly higher than PS2. Interestingly, the endogenous PS1:PS2 profile is maintained when PS is exogenously expressed. This is in contrast to the generally accepted rationale that ectopic gene expression using a constitutive promoter, such as CMV used in this study, will result in comparable protein levels of homologous proteins in the same cell lines. The differential PS1 and PS2 expression is likely the effect of post-translational protein regulation influenced by PS specific-localisation^31, 32, 57^ and the requisite involvement of other proteins for stable protein retention, namely Nicastrin, Aph1 and Pen-2.^1, 49–51, 58–60^ The transfection of exogenous PS into cells has shown that PS holoprotein is quickly degraded, while the endoproteolysed PS heterodimer fragments, by virtue of being incorporated into γ-secretase, are more stable.^61^ γ-Secretase incorporation of exogenously expressed PS is limited by the normal cellular regulation of the other complex components, and it appears from our data the innate cellular regulation of PS1 and PS2 is likely driven by the specific PS homologues. Equal ectopic expression of PS1- and PS2-γ-secretase could presumably be achieved by simultaneous overexpression of all γ-secretase components, such as that employed by Meckler and Checler.^31^ However, the extent to which PS expression is regulated by subcellular localisation, and organelle compartment size, remains unclear, and may be potentiated by the use of a variety of cell models.

Despite similar PS1:PS2 ratios, we show that variation in APP processing between the exogenous (ex) and endogenous (en) PS expression systems used in this study, prior to normalising for PS expression. A notable example of this is the difference in APP-CTF levels observed between expression systems (Fig 2, 5). Similarly, differences in the Aβ40 and Aβ42 levels generated between exogenous and endogenous systems results in different Aβ42:Aβ40 ratios between PS1 and PS2. Specifically, we observe that enPS2 generates a higher Aβ42:Aβ40 ratio, driven by lower Aβ40 production, compared with enPS1, while no difference is observed between exogenous PS. Our results for Notch1 processing, however, are recapitulated between the exogenous and endogenous system prior to normalisation of PS expression. There is no consensus in the literature regarding the Aβ42:Aβ40 ratio^30–32, 34, 36, 37^ or NICD^21, 34, 62 32, 62^ generated by PS1- vs PS2-γ-secretase, which is likely the result of the variety of expression systems, cell types and activity assays used. An additional difference observed between experimental systems in this study was that exPS1-NTF levels significantly increase in response to co-expression with either APP or Notch1, while exPS2-NTF levels do not differ. This effect is not observed in the endogenous PS system, and is further evidence that exogenous PS is not a faithful mimic of endogenous PS. The use of endogenous PS systems reduces the potential introduction of experimental artifacts and should be the preferred system, particularly given the current availability of CRISPR-Cas9 technology for cell line development.

Similar to other recent studies, we have used CRISPR-Cas9 technology to develop HEK-293 cell lines that retain enPS1 or enPS2 allowing us to study PS-specific γ-secretase function. While these recent studies use a variety of cell lines,^30, 32, 37^ the most comparable cell line to those used in this study are the HEK 293T cells developed by Lessard et al.^34^ These authors present both intracellular and extracellular levels of total Aβ, Aβ40 and Aβ42 using the HEK 293T cell line, and show that PS2 expressing cells generate significantly more intracellular total Aβ, Aβ40 and Aβ42.^34^ While we successfully measured extracellular Aβ, several attempts to detect intracellular Aβ were unsuccessful. We found that the amount of exogenous APP695Swe required for detection of intracellular Aβ by the ELISA assay used caused considerable cell death in the enPS1+ cells (Fig S5). This may be due to the use of the HEK 293 cell line in this study rather than HEK 293T used by Lessard et al. While we similarly show no significant difference in the absolute amount of secreted Aβ40 and Aβ42 between the PS1+ and PS2+ cell lines, we did show a significant increase in the Aβ42:Aβ40 ratio associated with enPS2 expression, due to accompanying reduction in Aβ40 generation. These findings are consistent with results reported in murine N2a cells lacking PS1^37^ and iPSC cells where PS1 and/or PS2 were conditionally knocked out and differentiated into neurons.^30^ Watanabe et al,^30^ further showed that during extended neuronal maturation of iPSCs there is a concomitant decrease in PS1 expression and increase in PS2 expression. Combined with our findings, these studies highlight the importance of PS2 in AD pathogenesis and the need to directly compare PS1 and PS2 expression to enable appropriate interpretation of PS specific γ-secretase contributions across multiple experimental systems.

There have been only a handful of studies that have considered PS expression, or mature nicastrin as a measure of γ-secretase expression. Yonemura and colleagues^41^ used myc-tagged exogenous PS in a yeast system, and demonstrate that PS1-NTF levels are approximately 28-times higher than PS2, and conclude that after normalising for expression there is no difference in overall activity. Lai et al,^44^ used radioactive labelling to determine PS1 and PS2 specific antibody sensitivity in order to calculate endogenous PS expression levels; in doing so, they identified that PS1 expression is approximately 1.4-times higher than PS2 in a murine blastocyte model. While Lai et al, ultimately concludes that PS1 generates more Aβ than PS2, they interestingly observe that while PS1 is similarly active in both membrane enriched cell-free γ-secretase assays and in-cell assays, PS2 is significantly less active in the membrane-based assay compared with the in-cell assay - an additional confounding feature to consider when interpreting the many γ-secretase activity studies. The use of mature nicastrin as a measure of active γ-secretase has also been used to normalise for activity between PS1 and PS2.^42, 43^ These studies conclude that PS1-γ-secretase generates more Aβ,^42, 43^ and similar levels of AICD and NICD;^42^ however, the use of cell-free membrane based assays,^43^ and poor evidence of nicastrin maturation,^42^ potentially confound interpretation. Our findings provide, to our knowledge, the first reported absolute quantitative measure of endogenous presenilin expression and demonstrate that PS2-γ-secretase processes equal amounts of Notch1 and more APP, compared to PS1-γ-secretase, when considered at an enzymatic unit level.

It should be acknowledged that our findings in HEK-293 cells cannot be generalised to other cell types which may show different presenilin expression profiles related to their developmental origin and function. Additionally, it must be noted that substrate specific localisation will likely effect the accessibility of PS1- and PS2-γ-secretase to certain substrate pools, influencing overall substrate processing capability.^30, 32^ However, we have developed useful tools that allow us to address this in future studies. Another potential limitation of this study is the use of incorporated PS expression as a measure of γ-secretase levels, which does not consider evidence suggesting that there is a pool of in-active γ-secretase. This evidence is from pulldown studies using modified γ-secretase inhibitors to capture ‘active’ γ-secretase complexes.^36, 63, 64^ The L685,458 inhibitor used as the basis for these studies does not however have equal affinity for PS1 and PS2.^13, 15^ Additionally, these experiments were performed using membrane extractions, which reduce PS2 - but not PS1 activity,^44^ potentially resulting in biased capture of PS1-complexes. Future experiments should consider the use of the BMS708163-derived γ-secretase capture tools,^65, 66^ which has comparable affinity for both PS1 and PS2.^6^ Additionally, these experiments should be developed in cell-based systems for accurate reflection of the active γ-secretase pool and can be used in conjunction with the PS quantitation method developed in this study for a more robust understanding of PS1 and PS2 specific γ-secretase expression and activity.

With the recognition that both PS1 and PS2 expression are important in the context of AD pathogenesis,^28, 39^ that PS1 and PS2 are differentially expressed expression throughout development,^30, 38, 40^ and varies between cell types,^32, 41, 44^ the field must look to methods that enable direct comparison of PS1 and PS2 expression when interpreting data. We acknowledge investigating individual endogenous PS in the absence of the homologous counterpart is not reflective of the physiological environment, and may belie the realities of the dynamic interplay between PS1 and PS2 for the other components of γ-secretase.^36, 67, 68^ However, we believe that coupling the use of an endogenous PS model, with a method to quantitate PS expression, presents a suitably representative experimental system to assess γ-secretase activity and the specific contributions of PS1 and PS2. To achieve this, we have developed and validated a PS1/2 fusion standard that, when used in conjunction with appropriate standard curves, calculates the number of PS protein units. This represents a novel approach to the absolute quantitation of presenilin levels in a cellular system that can be extended to tissues. Furthermore, we have demonstrated that endogenous PS expression is recapitulated in an exogenous expression system. While the resultant substrate processing is not in complete agreement, we have shown that PS2-γ-secretase is at least as active, if not more, than PS1-γ-secretase at processing both APP and Notch substrates.

There is a growing body of evidence that suggests that PS2 has greater implications for Aβ-related pathogenesis in AD then previously considered. In particular PS2-γ-secretase generates higher Aβ42:Aβ40 ratios shown in this study and others,^30, 36, 37^ and more intracellular Aβ.^32, 34^ Furthermore PS2 expression increases with neuronal maturation,^30, 69^ age, ^38, 40^ in response to PS1 mutations,^36^ and a rare autosomal dominant AD mutation.^28, 29^ We have presented here tools that enable accurate, direct comparison between PS1 and PS2 expression, and demonstrate how these can be used to improve our understanding and interpretation of the effect of PS expression on γ-secretase activity.

## DATA AVAILABILITY

The data that support the findings of this study are available in the methods and/or supplementary material of this article.

## AUTHOR CONTRIBUTIONS

M. Eccles and G. Verdile conceived and designed the research. M. Eccles, N. Main, M. Sabale and B. Roberts-Mok performed the experiments and acquired the data. M. Eccles, P.E. Fraser and G. Verdile analysed and interpreted data. All authors were involved in manuscript preparation.

## ACKNOWLEDGEMENTS

M. Eccles is supported by Dementia Australia Research Foundation.

## REFERENCES

1. Kimberly, W. T., LaVoie, M. J., Ostaszewski, B. L., Ye, W., Wolfe, M. S., and Selkoe, D. J. (2003) γ-Secretase is a membrane protein complex comprised of presenilin, nicastrin, aph-1, and pen-2. Proceedings of the National Academy of Sciences 100, 6382–6387

2. Güner, G., and Lichtenthaler, S. F. (2020) The substrate repertoire of γ-secretase/presenilin. Seminars in Cell & Developmental Biology 105, 27–42

3. Kimberly, W. T., Xia, W., Rahmati, T., Wolfe, M. S., and Selkoe, D. J. (2000) The transmembrane aspartates in presenilin 1 and 2 are obligatory for γ-secretase activity and amyloid β-protein generation. Journal of Biological Chemistry 275, 3173–3178

4. De Strooper, B., Annaert, W., Cupers, P., Saftig, P., Craessaerts, K., Mumm, J. S., Schroeter, E. H., Schrijvers, V., Wolfe, M. S., Ray, W. J., Goate, A., and Kopan, R. (1999) A presenilin-1-dependent [gamma]-secretase-like protease mediates release of Notch intracellular domain. Nature 398, 518–522

5. DeTure, M. A., and Dickson, D. W. (2019) The neuropathological diagnosis of Alzheimer’s disease. Molecular Neurodegeneration 14, 32

6. Borgegård, T., Gustavsson, S., Nilsson, C., Parpal, S., Klintenberg, R., Berg, A.-L., Rosqvist, S., Serneels, L., Svensson, S., Olsson, F., Jin, S., Yan, H., Wanngren, J., Jureus, A., Ridderstad-Wollberg, A., Wollberg, P., Stockling, K., Karlström, H., Malmberg, Å., Lund, J., Arvidsson, P. I., De Strooper, B., Lendahl, U., and Lundkvist, J. (2012) Alzheimer’s disease: Presenilin-2 sparing γ-secretase inhibition is a tolerable Aβ peptide-lowering strategy. The Journal of Neuroscience 32, 17297–17305

7. Doody, R. S., Raman, R., Farlow, M., Iwatsubo, T., Vellas, B., Joffe, S., Kieburtz, K., He, F., Sun, X., Thomas, R. G., Aisen, P. S., Siemers, E., Sethuraman, G., and Mohs, R. (2013) A phase 3 trial of semagacestat for treatment of Alzheimer’s disease. New England Journal of Medicine 369, 341–350

8. Gillman, K. W., Starrett, J. E., Parker, M. F., Xie, K., Bronson, J. J., Marcin, L. R., McElhone, K. E., Bergstrom, C. P., Mate, R. A., Williams, R., Meredith, J. E., Burton, C. R., Barten, D. M., Toyn, J. H., Roberts, S. B., Lentz, K. A., Houston, J. G., Zaczek, R., Albright, C. F., Decicco, C. P., Macor, J. E., and Olson, R. E. (2010) Discovery and evaluation of BMS-708163, a potent, selective and orally bioavailable γ-secretase inhibitor. ACS Medicinal Chemistry Letters 1, 120–124

9. Golde, T. E., Koo, E. H., Felsenstein, K. M., Osborne, B. A., and Miele, L. (2013) γ-Secretase inhibitors and modulators. Biochimica et Biophysica Acta (BBA) - Biomembranes 1828, 2898–2907

10. Shearman, M. S., Beher, D., Clarke, E. E., Lewis, H. D., Harrison, T., Hunt, P., Nadin, A., Smith, A. L., Stevenson, G., and Castro, J. L. (2000) L-685,458, an aspartyl protease transition state mimic, is a potent inhibitor of amyloid beta-protein precursor gamma-secretase activity. Biochemistry 39, 8698–8704

11. Coric, V., van Dyck, C. H., Salloway, S., and, et al. (2012) Safety and tolerability of the γ-secretase inhibitor avagacestat in a phase 2 study of mild to moderate Alzheimer disease. Archives of Neurology 69, 1430–1440

12. Yang, Z. Y., Li, J. M., Xiao, L., Mou, L., Cai, Y., Huang, H., Luo, X. G., and Yan, X. X. (2014) [(3) H]-L685,458 binding sites are abundant in multiple peripheral organs in rats: implications for safety assessment of putative γ-secretase targeting drugs. Basic & Clinical Pharmacology & Toxicology 115, 518–526

13. Ebke, A., Luebbers, T., Fukumori, A., Shirotani, K., Haass, C., Baumann, K., and Steiner, H. (2011) Novel γ-secretase enzyme modulators directly target presenilin protein. Journal of Biological Chemistry 286, 37181–37186

14. Guo, X., Wang, Y., Zhou, J., Jin, C., Wang, J., Jia, B., Jing, D., Yan, C., Lei, J., Zhou, R., and Shi, Y. (2022) Molecular basis for isoform-selective inhibition of presenilin-1 by MRK-560. Nature Communications 13, 6299

15. Lee, J., Song, L., Terracina, G., Bara, T., Josien, H., Asberom, T., Sasikumar, T. K., Burnett, D. A., Clader, J., Parker, E. M., and Zhang, L. (2011) Identification of presenilin 1-selective γ-secretase inhibitors with reconstituted γ-secretase complexes. Biochemistry 50, 4973–4980

16. Bursavich, M. G., Harrison, B. A., and Blain, J.-F. (2016) Gamma secretase modulators: New Alzheimer’s drugs on the horizon? Journal of Medicinal Chemistry 59, 7389–7409

17. Oehlrich, D., Berthelot, D. J., and Gijsen, H. J. (2011) γ-Secretase modulators as potential disease modifying anti-Alzheimer’s drugs. Journal of Medicinal Chemistry 54, 669–698

18. Kounnas, M. Z., Lane-Donovan, C., Nowakowski, D. W., Herz, J., and Comer, W. T. (2017) NGP 555, a γ-secretase modulator, lowers the amyloid biomarker, Aβ42, in cerebrospinal fluid while preventing Alzheimer’s disease cognitive decline in rodents. Alzheimer’s & Dementia: Translational Research & Clinical Interventions 3, 65–73

19. Mumm, J. S., Schroeter, E. H., Saxena, M. T., Griesemer, A., Tian, X., Pan, D. J., Ray, W. J., and Kopan, R. (2000) A ligand-induced extracellular cleavage regulates γ-secretase-like proteolytic activation of notch1. Molecular Cell 5, 197–206

20. Buxbaum, J. D., Liu, K. N., Luo, Y., Slack, J. L., Stocking, K. L., Peschon, J. J., Johnson, R. S., Castner, B. J., Cerretti, D. P., and Black, R. A. (1998) Evidence that tumor necrosis factor alpha converting enzyme is involved in regulated alpha-secretase cleavage of the Alzheimer amyloid protein precursor. Journal of Biological Chemistry 273, 27765–27767

21. Zhang, Z., Nadeau, P., Song, W., Donoviel, D., Yuan, M., Bernstein, A., and Yankner, B. A. (2000) Presenilins are required for [gamma]-secretase cleavage of [beta]-APP and transmembrane cleavage of Notch-1. Nature Cell Biology 2, 463–465

22. Lammich, S., Kojro, E., Postina, R., Gilbert, S., Pfeiffer, R., Jasionowski, M., Haass, C., and Fahrenholz, F. (1999) Constitutive and regulated alpha-secretase cleavage of Alzheimer’s amyloid precursor protein by a disintegrin metalloprotease. Proceedings of the National Academy of Sciences 96, 3922–3927

23. Vassar, R., Bennett, B. D., Babu-Khan, S., Kahn, S., Mendiaz, E. A., Denis, P., Teplow, D. B., Ross, S., Amarante, P., Loeloff, R., Luo, Y., Fisher, S., Fuller, J., Edenson, S., Lile, J., Jarosinski, M. A., Biere, A. L., Curran, E., Burgess, T., Louis, J.-C., Collins, F., Treanor, J., Rogers, G., and Citron, M. (1999) β-Secretase Cleavage of Alzheimer’s Amyloid Precursor Protein by the Transmembrane Aspartic Protease BACE. Science 286, 735–741

24. Takami, M., Nagashima, Y., Sano, Y., Ishihara, S., Morishima-Kawashima, M., Funamoto, S., and Ihara, Y. (2009) γ-Secretase: Successive tripeptide and tetrapeptide release from the transmembrane domain of β-carboxyl terminal fragment. The Journal of Neuroscience 29, 13042–13052

25. Matsumura, N., Takami, M., Okochi, M., Wada-Kakuda, S., Fujiwara, H., Tagami, S., Funamoto, S., Ihara, Y., and Morishima-Kawashima, M. (2014) γ-Secretase associated with lipid rafts: multiple interactive pathways in the stepwise processing of β-carboxyl-terminal fragment. Journal of Biological Chemistry 289, 5109–5121

26. Alzforum. Mutation Database. https://www.alzforum.org/mutations. Accessed September 15, 2022.

27. Ryman, D. C., Acosta-Baena, N., Aisen, P. S., Bird, T., Danek, A., Fox, N. C., Goate, A., Frommelt, P., Ghetti, B., Langbaum, J. B. S., Lopera, F., Martins, R., Masters, C. L., Mayeux, R. P., McDade, E., Moreno, S., Reiman, E. M., Ringman, J. M., Salloway, S., Schofield, P. R., Sperling, R., Tariot, P. N., Xiong, C., Morris, J. C., Bateman, R. J., and And the Dominantly Inherited Alzheimer, N. (2014) Symptom onset in autosomal dominant Alzheimer disease: A systematic review and meta-analysis. Neurology 83, 253–260

28. Pang, Y., Li, T., Wang, Q., Qin, W., Li, Y., Wei, Y., and Jia, L. (2021) A rare variation in the 3’ untranslated region of the presenilin 2 gene is linked to Alzheimer’s disease. Molecular Neurobiology 58, 4337–4347

29. Jia, L., Fu, Y., Shen, L., Zhang, H., Zhu, M., Qiu, Q., Wang, Q., Yan, X., Kong, C., Hao, J., Wei, C., Tang, Y., Qin, W., Li, Y., Wang, F., Guo, D., Zhou, A., Zuo, X., Yu, Y., Li, D., Zhao, L., Jin, H., and Jia, J. (2020) PSEN1, PSEN2, and APP mutations in 404 Chinese pedigrees with familial Alzheimer’s disease. Alzheimer’s & Dementia 16, 178–191

30. Watanabe, H., Imaizumi, K., Cai, T., Zhou, Z., Tomita, T., and Okano, H. (2021) Flexible and accurate substrate processing with distinct presenilin/γ-secretases in human cortical neurons. eNeuro 8, ENEURO.0500-0520.2021

31. Meckler, X., and Checler, F. (2016) Presenilin 1 and presenilin 2 target γ-secretase complexes to distinct cellular compartments. Journal of Biological Chemistry 291, 12821–12837

32. Sannerud, R., Esselens, C., Ejsmont, P., Mattera, R., Rochin, L., Tharkeshwar, Arun K., De Baets, G., De Wever, V., Habets, R., Baert, V., Vermeire, W., Michiels, C., Groot, Arjan J., Wouters, R., Dillen, K., Vints, K., Baatsen, P., Munck, S., Derua, R., Waelkens, E., Basi, Guriqbal S., Mercken, M., Vooijs, M., Bollen, M., Schymkowitz, J., Rousseau, F., Bonifacino, Juan S., Van Niel, G., De Strooper, B., and Annaert, W. (2016) Restricted location of PSEN2/γ-secretase determines substrate specificity and generates an intracellular Aβ pool. Cell 166, 193–208

33. Zhu, L., Su, M., Lucast, L., Liu, L., Netzer, W. J., Gandy, S. E., and Cai, D. (2012) Dynamin 1 regulates amyloid generation through modulation of BACE-1. PLoS One 7, e45033

34. Lessard, C. B., Rodriguez, E., Ladd, T. B., Minter, L. M., Osborne, B. A., Miele, L., Golde, T. E., and Ran, Y. (2019) Individual and combined presenilin 1 and 2 knockouts reveal that both have highly overlapping functions in HEK293T cells. Journal of Biological Chemistry

35. Acx, H., Chávez-Gutiérrez, L., Serneels, L., Lismont, S., Benurwar, M., Elad, N., and De Strooper, B. (2014) Signature amyloid β profiles are produced by different γ-secretase complexes. Journal of Biological Chemistry 289, 4346–4355

36. Placanica, L., Tarassishin, L., Yang, G., Peethumnongsin, E., Kim, S.-H., Zheng, H., Sisodia, S. S., and Li, Y.-M. (2009) Pen2 and presenilin-1 modulate the dynamic equilibrium of presenilin-1 and presenilin-2 γ-secretase complexes. Journal of Biological Chemistry 284, 2967–2977

37. Pimenova, A. A., and Goate, A. M. (2020) Novel presenilin 1 and 2 double knock-out cell line for in vitro validation of PSEN1 and PSEN2 mutations. Neurobiology of Disease, 104785

38. Lee, M. K., Slunt, H. H., Martin, L. J., Thinakaran, G., Kim, G., Gandy, S. E., Seeger, M., Koo, E., Price, D. L., and Sisodia, S. S. (1996) Expression of presenilin 1 and 2 (PS1 and PS2) in human and murine tissues. The Journal of Neuroscience 16, 7513–7525

39. Davidsson, P., Bogdanovic, N., Lannfelt, L., and Blennow, K. (2001) Reduced expression of amyloid precursor protein, presenilin-1 and rab3a in cortical brain regions in Alzheimer’s disease. Dementia and Geriatric Cognitive Disorders 12, 243–250

40. Thakur, M. K., and Ghosh, S. (2007) Age and sex dependent alteration in presenilin expression in mouse cerebral cortex. Cellular and Molecular Neurobiology 27, 1059–1067

41. Yonemura, Y., Futai, E., Yagishita, S., Suo, S., Tomita, T., Iwatsubo, T., and Ishiura, S. (2011) Comparison of presenilin 1 and presenilin 2 γ-secretase activities using a yeast reconstitution system. Journal of Biological Chemistry 286, 44569–44575

42. Pintchovski, S. A., Schenk, D. B., and Basi, G. S. (2013) Evidence that enzyme processivity mediates differential Aβ production by PS1 and PS2. Current Alzheimer Research 10, 4–10

43. Shirotani, K., Tomioka, M., Kremmer, E., Haass, C., and Steiner, H. (2007) Pathological activity of familial Alzheimer’s disease-associated mutant presenilin can be executed by six different γ-secretase complexes. Neurobiology of Disease 27, 102–107

44. Lai, M.-T., Chen, E., Crouthamel, M.-C., DiMuzio-Mower, J., Xu, M., Huang, Q., Price, E., Register, R. B., Shi, X.-P., Donoviel, D. B., Bernstein, A., Hazuda, D., Gardell, S. J., and Li, Y.-M. (2003) Presenilin-1 and presenilin-2 exhibit distinct yet overlapping γ-secretase activities. Journal of Biological Chemistry 278, 22475–22481

45. Ran, F. A., Hsu, P. D., Wright, J., Agarwala, V., Scott, D. A., and Zhang, F. (2013) Genome engineering using the CRISPR-Cas9 system. Nature Protocols 8, 2281

46. Ye, J., Coulouris, G., Zaretskaya, I., Cutcutache, I., Rozen, S., and Madden, T. L. (2012) Primer-BLAST: a tool to design target-specific primers for polymerase chain reaction. BMC Bioinformatics 13, 134

47. Koch, P., Tamboli, I. Y., Mertens, J., Wunderlich, P., Ladewig, J., Stüber, K., Esselmann, H., Wiltfang, J., Brüstle, O., and Walter, J. (2012) Presenilin-1 L166P mutant human pluripotent stem cell-derived neurons exhibit partial loss of γ-secretase activity in endogenous amyloid-β generation. The American Journal of Pathology 180, 2404–2416

48. Pfaffl, M. W. (2001) A new mathematical model for relative quantification in real-time RT-PCR. Nucleic acids research 29, e45–e45

49. Steiner, H., Winkler, E., Edbauer, D., Prokop, S., Basset, G., Yamasaki, A., Kostka, M., and Haass, C. (2002) PEN-2 is an integral component of the γ-secretase complex required for coordinated expression of presenilin and nicastrin. Journal of Biological Chemistry 277, 39062–39065

50. Yang, D.-S., Tandon, A., Chen, F., Yu, G., Yu, H., Arawaka, S., Hasegawa, H., Duthie, M., Schmidt, S. D., Ramabhadran, T. V., Nixon, R. A., Mathews, P. M., Gandy, S. E., Mount, H. T. J., St George-Hyslop, P., and Fraser, P. E. (2002) Mature glycosylation and trafficking of nicastrin modulate its binding to presenilins. Journal of Biological Chemistry 277, 28135–28142

51. Luo, W. J., Wang, H., Li, H., Kim, B. S., Shah, S., Lee, H. J., Thinakaran, G., Kim, T. W., Yu, G., and Xu, H. (2003) PEN-2 and APH-1 coordinately regulate proteolytic processing of presenilin 1. Journal of Biological Chemistry 278, 7850–7854

52. Bai, X.-c., Yan, C., Yang, G., Lu, P., Ma, D., Sun, L., Zhou, R., Scheres, S. H. W., and Shi, Y. (2015) An atomic structure of human γ-secretase. Nature 525, 212–217

53. Guan, Y., Zhu, Q., Huang, D., Zhao, S., Jan Lo, L., and Peng, J. (2015) An equation to estimate the difference between theoretically predicted and SDS PAGE-displayed molecular weights for an acidic peptide. Scientific Reports 5, 13370

54. Tiwari, P., Kaila, P., and Guptasarma, P. (2019) Understanding anomalous mobility of proteins on SDS-PAGE with special reference to the highly acidic extracellular domains of human E- and N-cadherins. Electrophoresis 40, 1273–1281

55. Sato, T., Diehl, T. S., Narayanan, S., Funamoto, S., Ihara, Y., De Strooper, B., Steiner, H., Haass, C., and Wolfe, M. S. (2007) Active γ-secretase complexes contain only one of each component. Journal of Biological Chemistry 282, 33985–33993

56. Thinakaran, G., Borchelt, D. R., Lee, M. K., Slunt, H. H., Spitzer, L., Kim, G., Ratovitsky, T., Davenport, F., Nordstedt, C., Seeger, M., Hardy, J., Levey, A. I., Gandy, S. E., Jenkins, N. A., Copeland, N. G., Price, D. L., and Sisodia, S. S. (1996) Endoproteolysis of presenilin 1 and accumulation of processed derivatives in vivo. Neuron 17, 181–190

57. Yousefi, R., Jevdokimenko, K., Kluever, V., Pacheu-Grau, D., and Fornasiero, E. F. (2021) Influence of subcellular localization and functional state on protein turnover. Cells 10

58. Fraser, P. E., Levesque, G., Yu, G., Mills, L. R., Thirlwell, J., Frantseva, M., Gandy, S. E., Seeger, M., Carlen, P. L., and St George-Hyslop, P. (1998) Presenilin 1 is actively degraded by the 26S proteasome. Neurobiology of Aging 19, S19–S21

59. Edbauer, D., Winkler, E., Haass, C., and Steiner, H. (2002) Presenilin and nicastrin regulate each other and determine amyloid β-peptide production via complex formation. Proceedings of the National Academy of Sciences 99, 8666–8671

60. Thinakaran, G., Harris, C. L., Ratovitski, T., Davenport, F., Slunt, H. H., Price, D. L., Borchelt, D. R., and Sisodia, S. S. (1997) Evidence that levels of presenilins (PS1 and PS2) are coordinately regulated by competition for limiting cellular factors. Journal of Biological Chemistry 272, 28415–28422

61. Zhang, J., Kang, D. E., Xia, W., Okochi, M., Mori, H., Selkoe, D. J., and Koo, E. H. (1998) Subcellular distribution and turnover of presenilins in transfected cells. Journal of Biological Chemistry 273, 12436–12442

62. Yonemura, Y., Futai, E., Yagishita, S., Kaether, C., and Ishiura, S. (2016) Specific combinations of presenilins and Aph1s affect the substrate specificity and activity of γ-secretase. Biochemical and Biophysical Research Communications 478, 1751–1757

63. Frånberg, J., Svensson, A. I., Winblad, B., Karlström, H., and Frykman, S. (2011) Minor contribution of presenilin 2 for γ-secretase activity in mouse embryonic fibroblasts and adult mouse brain. Biochemical and Biophysical Research Communications 404, 564–568

64. Teranishi, Y., Hur, J.-Y., Welander, H., Frånberg, J., Aoki, M., Winblad, B., Frykman, S., and Tjernberg, L. O. (2010) Affinity pulldown of γ-secretase and associated proteins from human and rat brain. Journal of Cellular and Molecular Medicine 14, 2675–2686

65. Crump, C. J., Murrey, H. E., Ballard, T. E., Am Ende, C. W., Wu, X., Gertsik, N., Johnson, D. S., and Li, Y. M. (2016) Development of sulfonamide photoaffinity inhibitors for probing cellular γ-secretase. ACS Chemical Neuroscience 7, 1166–1173

66. Gertsik, N., am Ende, C. W., Geoghegan, K. F., Nguyen, C., Mukherjee, P., Mente, S., Seneviratne, U., Johnson, D. S., and Li, Y.-M. (2017) Mapping the binding site of BMS-708163 on γ-secretase with cleavable photoprobes. Cell Chemical Biology 24, 3–8

67. Chen, F., Tandon, A., Sanjo, N., Gu, Y.-J., Hasegawa, H., Arawaka, S., Lee, F. J. S., Ruan, X., Mastrangelo, P., Erdebil, S., Wang, L., Westaway, D., Mount, H. T. J., Yankner, B., Fraser, P. E., and George-Hyslop, P. S. (2003) Presenilin 1 and presenilin 2 have differential effects on the stability and maturation of nicastrin in mammalian brain. Journal of Biological Chemistry 278, 19974–19979

68. Pardossi-Piquard, R., Yang, S. P., Kanemoto, S., Gu, Y., Chen, F., Böhm, C., Sevalle, J., Li, T., Wong, P. C., Checler, F., Schmitt-Ulms, G., St. George-Hyslop, P., and Fraser, P. E. (2009) APH1 polar transmembrane residues regulate the assembly and activity of presenilin complexes. Journal of Biological Chemistry 284, 16298–16307

69. Culvenor, J. G., Evin, G., Cooney, M. A., Wardan, H., Sharples, R. A., Maher, F., Reed, G., Diehlmann, A., Weidemann, A., Beyreuther, K., and Masters, C. L. (2000) Presenilin 2 expression in neuronal cells: induction during differentiation of embryonic carcinoma cells. Experimental Cell Research 255, 192–206

