## Supplemental Data for "Quantitative Comparison of Presenilin Protein Expression Reveals Greater Activity of PS2-γ-Secretase"

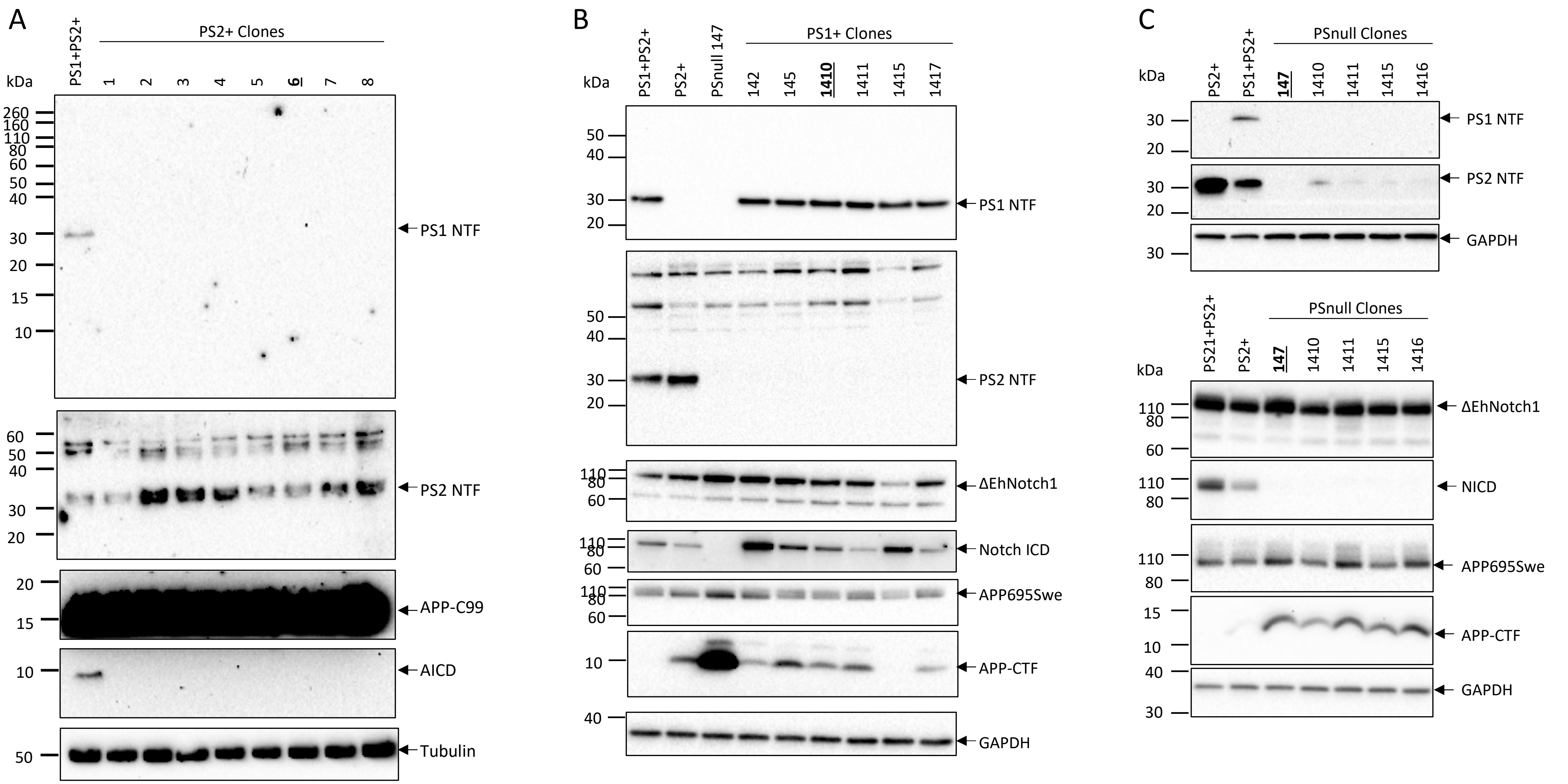


**Figure S1** Multiple clones of presenilin knockout cell lines were assessed for PS1 and PS2 expression and either AICD generation via cell-free assay using crude CHAPSO cell membrane lysates, or APP-CTF accumulation and NICD generation with hAPP695Swe or ΔEhNotch1 transient transfection. Clones were selected on the bases of complete loss of appropriate PS homologues and average expression or processing compared to other clones. Immunoblot results for **A)** PS2+ clones assessed for PS1 and PS2 NTF expression and AICD generation. **B)** PS1+ clones assessed for PS1 and PS2 NTF expression, APP-CTF accumulation and NICD generation **C)** PSnull clones assessed for PS1 and PS2 NTF expression, APP-CTF accumulation and NICD generation. Selected clone numbers are bolded and underlined.

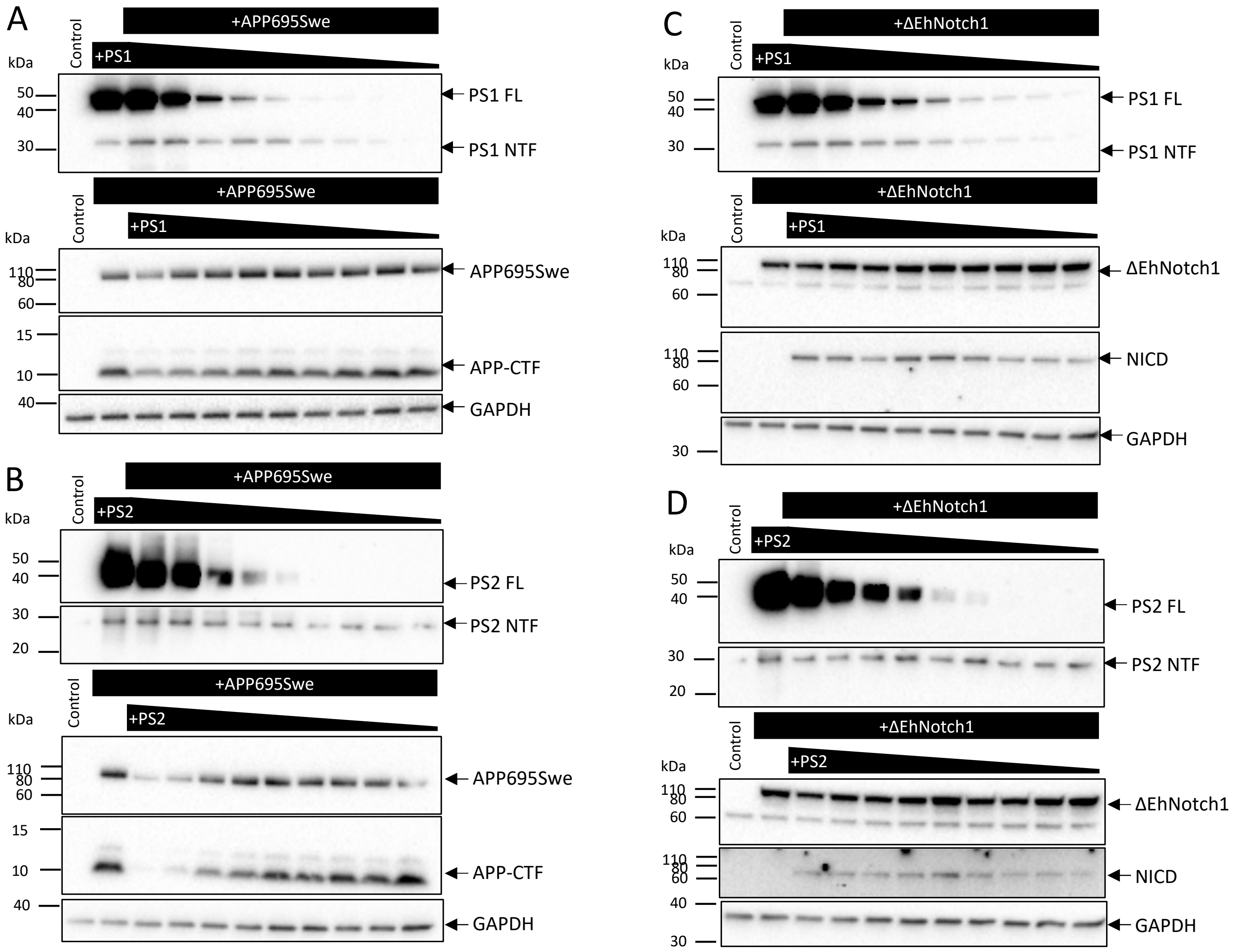


**Figure S2** Titration of exogeneous PS1 and PS2 transfections to determine amount of PS that maximises PS-NTF incorporation and minimises the amount of PS-FL overexpression. The selected amount of PS used for experiments was 116ng for transfection with APP695Swe and 111ng for transfection with ΔEhNotch1 (Note: Substrate was transfected at 3:1 Substrate-vector:PS-vector ratio, all ng amounts were calculated based on specific vector base pair sizes) **A)** PS1 co-transfected with APP695Swe **B)** PS2 co-transfected with APP695Swe **C)** PS1 co-transfected with ΔEhNotch1 **D )** PS2 co-transfected with ΔEhNotch1.

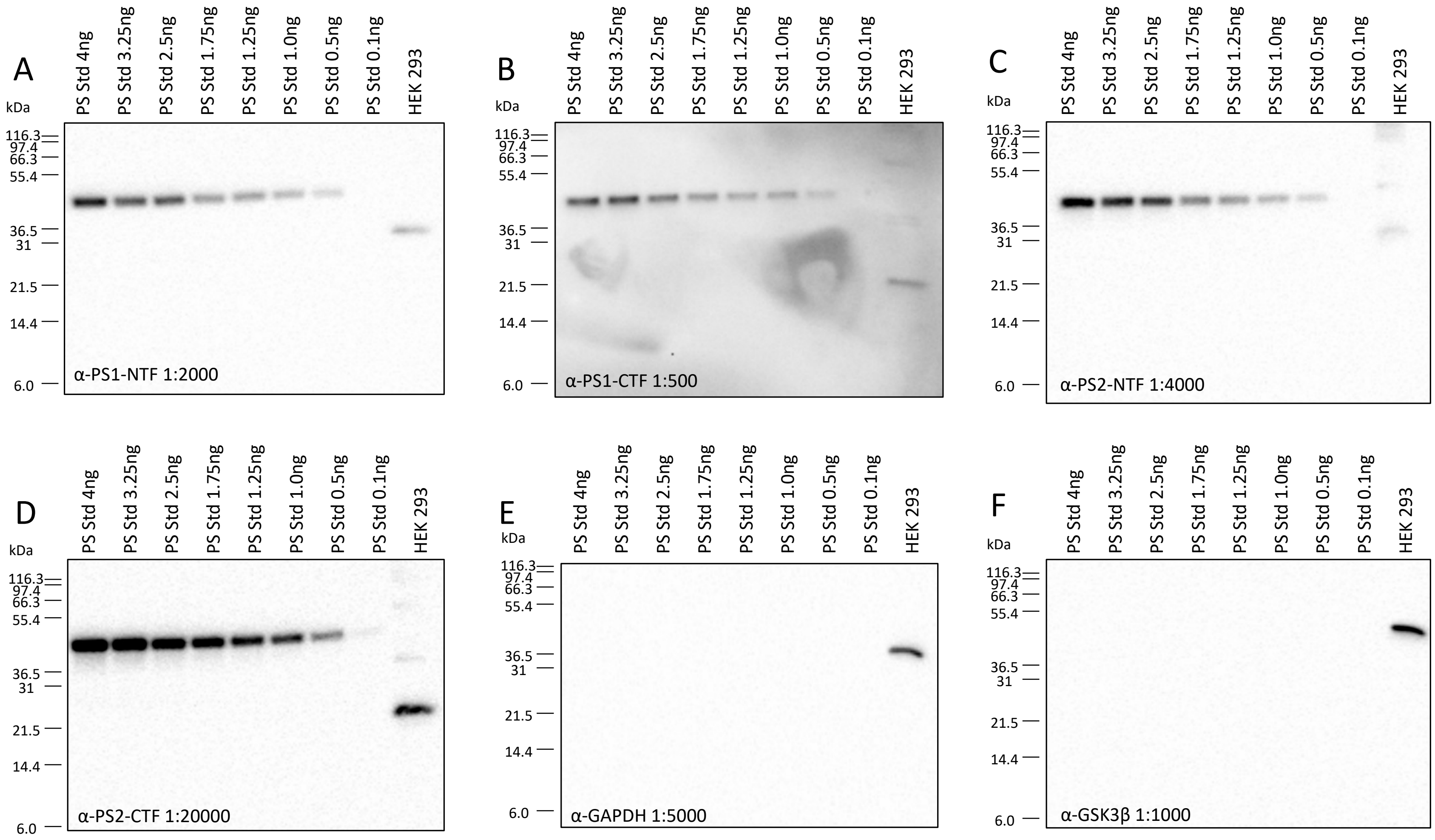


**Figure S3** PS-std incorporates N-terminal and cytoplasmic loop antibody epitope regions of human PS1 and PS2 and is detected by multiple commercial PS antibodies, but not by antibodies targeting unrelated proteins. **A)** PS1 NTF – Biolegend (823401) **B)** PS1 CTF – Abcam (ab15458) **C)** PS2 NTF in house antibody courtesy P.E. Fraser **D)** PS2 CTF – Abcam (ab51249) **D)** GAPDH – Cell Signalling (5174) **E)** GSK3β (D5C5Z) – Cell Signalling (12456)

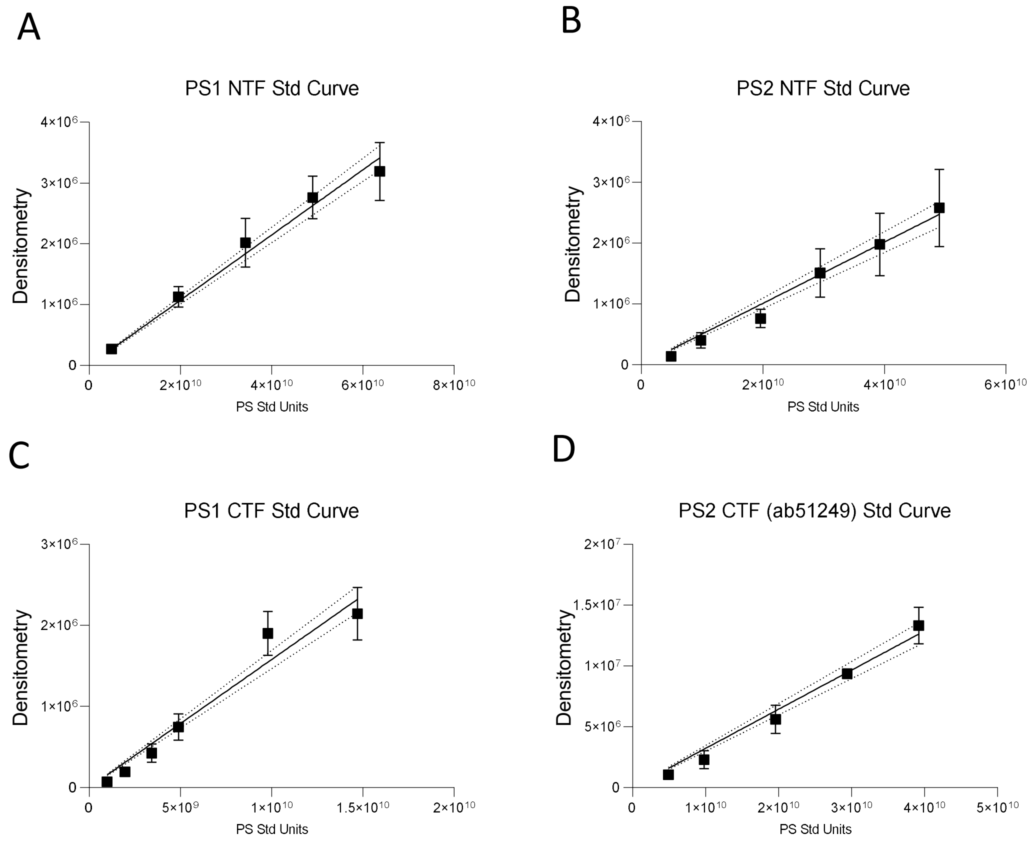


**Figure S4** Endogenous PS quantitation standard curves generated by plotting the number of PS-Std Units loaded against the quantitated band densitometry to determine equation then applied to the band densitometry for the endogenous PS detected in whole cell lysates. Standard curves shown here for quantitation shown in Figure 4. Standard curves are specific to the antibody used and the replicate set of immunoblots. **A)** PS1 NTF – Biolegend (823401) **B)** PS2 NTF – Biolegend (814204) **C)** PS1 CTF – Cell Signalling (5643S) **D)** PS2 CTF – Abcam (ab51249)

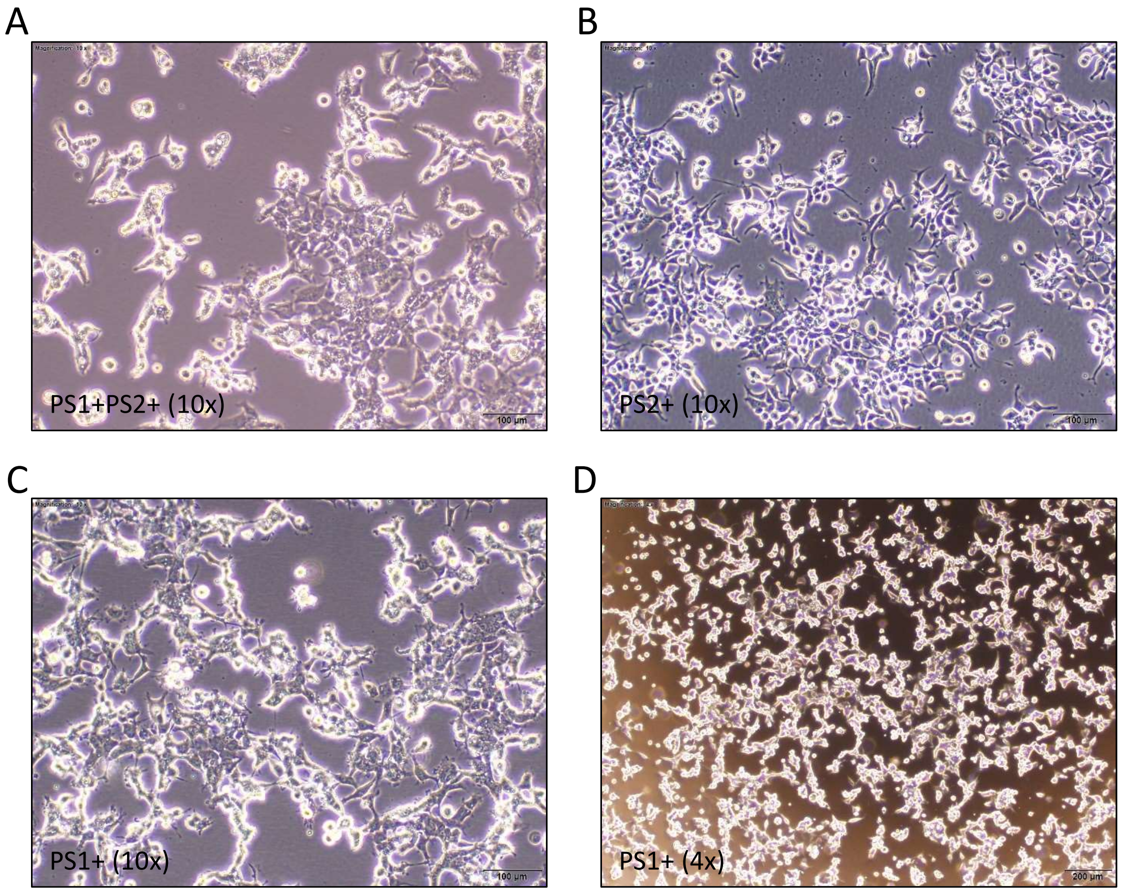


**Figure S5** Cell line transfection of hAPP695Swe for intracellular Aβ detection caused cell death in PS1+ cells. The amount of hAPP695Swe construct required to detect intracellular Aβ using either human Aβ40 ELISA (Thermofisher KHB3482) or human Aβ42 Ultrasensitive ELISA (Thermofisher KHB3544) had to be increased to 2500 ng per 6 well dish compared with 384 ng used for detection in conditioned media. Significant cell death was observed, particularly in the PS1+ cells, making the experiment untenable. Bright field microscopy images of cells approximately 20hrs post transfection **A)** PS1+PS2+ cells at 10x magnification **B)** PS2+ cells at 10x magnification **C-D)** PS1+ cells at 10x and 4x magnification respectively.

**Table S3 List of Commercial Antibodies for potential use with PS-Std**

| Presenilin | Fragment | Clone | Supplier & Catalogue# | | Epitope region (aa) | In PS Std | Tested with PS Std |
|---|---|---|---|---|---|---|---|
| **PS1** | **NTF** | **NT1** | **Biolegend** | **823401** | **41-49** | **YES** | **YES** |
| PS1 | NTF | | Abcam | ab252856 | proprietry | Unknown | No |
| PS1 | NTF | | Abcam | ab216400 | 1-100 | Partial | No |
| PS1 | NTF | | Abcam | ab255850 | proprietry | Unknown | No |
| PS1 | NTF | APS11 | Multiple | | 21-34 | YES | No |
| PS1 | NTF | | ThermoFisher | PA5-98093 | 1-160 | Partial | No |
| PS1 | NTF | ARC0440 | ThermoFisher | MA5-35263 | 1-100 | Partial | No |
| PS1 | NTF | | ThermoFisher | PA5-119872 | 330-380 | YES | No |
| PS1 | NTF | | ThermoFisher | PA5-98092 | 4-218 | Partial | No |
| PS1 | NTF | | ThermoFisher/Fabgennix | PSEN-101AP | 1-50 | Partial | No |
| PS1 | NTF | | Sigma-Aldrich | MAB1563 | 21-80 | Partial | No |
| PS1 | NTF | | ABclonal | A19103 | 1-100 | Partial | No |
| **PS1** | **CTF** | **D39D1** | **Cell Signalling** | **5643S** | **300-380** | **YES** | **YES** |
| PS1 | CTF | | Cell Signalling | 3622 | around V293 | YES | No |
| **PS1** | **CTF** | **APS18** | **Multiple** | | **313-334** | **Partial** | **YES** |
| PS1 | CTF | | Abcam | EP2000Y | proprietry | Unknown | No |
| PS1 | CTF | | ThermoFisher | BS-0025M | 299-350 | YES | No |
| PS1 | CTF | | ThermoFisher | PA5-96088 | 290-380 | Partial | No |
| PS1 | CTF | | ThermoFisher | PA5-30585 | 257-352 | Partial | No |
| PS1 | CTF | | Sigma-Aldrich | AB5308 | 275-367 | Partial | No |
| PS1 | CTF | | Sigma-Aldrich | MAB5232 | 263-378 | Partial | No |
| PS1 | CTF | | ABclonal | A2187 | 290-380 | YES | No |
| **PS2** | **NTF** | | **Biolegend** | **814204** | **1-87** | **YES** | **YES** |
| **PS2** | **NTF** | **APS21** | **Multiple** | | **31-45** | **YES** | **YES** |
| PS2 | NTF | | ThermoFisher | PA5-112675 | 7-77 | YES | No |
| PS2 | NTF | | ThermoFisher | PA5-94969 | 39-51 | YES | No |
| PS2 | NTF | | ThermoFisher/Bioss | BS-3815R | 51-150 | Partial | No |
| PS2 | NTF | | Santa-Cruz | sc-393758 | 1-76 | YES | No |
| PS2 | NTF | | ABclonal | A7719 | 1-75 | YES | No |
| **PS2** | **CTF** | **EP1515Y** | **Abcam** | **ab51249** | **300-400** | **Partial** | **YES** |
| PS2 | CTF | | ThermoFisher/Bethyl Laboratories | A304-342A | 275-325 | Partial | No |
| PS2 | CTF | | ThermoFisher | PA5-96774 | 300-400 | Partial | No |
| PS2 | CTF | | ThermoFisher | PA5-115793 | 316-344 | YES | No |
